# ARCADIA Reveals Spatially Dependent Transcriptional Programs through Integration of scRNA-seq and Spatial Proteomics

**DOI:** 10.1101/2025.11.20.689521

**Authors:** Bar Rozenman, Kevin Hoffer-Hawlik, Nicholas Djedjos, Elham Azizi

**Author notes:** Equal contribution (Bar Rozenman, Kevin Hoffer-Hawlik).

## Abstract

**Motivation:** Cellular states are strongly influenced by spatial context, but single-cell RNA sequencing (scRNA-seq) loses information about local tissue organization, while spatial proteomic assays capture limited marker panels that constrain transcriptomic inference. Integrating these modalities can elucidate how spatial niches shape transcriptional programs, yet existing approaches depend on either feature-level correspondence such as gene–protein linkage or cell-level barcode pairing, which is often unavailable.

**Results:** We present ARCADIA (ARchetype-based Clustering and Alignment with Dual Integrative Autoencoders), a generative framework for cross-modal integration that operates without cell barcode pairing and does not assume direct feature-to-feature correspondence. ARCADIA identifies modality-specific archetypes, i.e., convex combinations of cells representing extreme phenotypic states, and aligns these anchors across modalities by minimizing the discrepancy between their cell-type composition profiles. The aligned archetypes define a shared coordinate system that anchors dual variational autoencoders (VAEs) trained with cross-modal geometric regularization, preserving archetype structure and spatial neighborhood information while enabling bidirectional translation between modalities. On semi-synthetic CITE-seq data, ARCADIA outperforms existing weak-linkage methods. Applied to independent human tonsil scRNA-seq and CODEX data, ARCADIA reconstructs known tissue architecture and reveals spatially dependent transcriptional programs linking B-cell maturation and T-cell activation or exhaustion to microenvironmental niches.

**Availability and Implementation:** Source code is accessible at https://github.com/azizilab/ARCADIA_public. Reproducibility scripts and data are available at https://github.com/azizilab/arcadia_reproducibility.

## Introduction

Cell function is strongly influenced by spatial context in tissue. Studying spatial organization and cell-cell communication with transcriptomics reveals coordinated molecular networks of development, disease evolution, therapy response, and microenvironment adaptation [1, 2, 3]. Single-cell RNA sequencing (scRNA-seq) enables high-resolution profiling of cell phenotypes and states, yet dissociation removes spatial context [4, 5]. Spatial assays including spatial transcriptomics (ST) give insight into tissue organization, but ST is limited in unbiased discovery of gene programs and signaling pathways within neighborhoods, as it sacrifices throughput (e.g. limited panels in Xenium, merFISH) or resolution (e.g. multi-cell spots in Visium, Slide-seq; cross-spot diffusion in VisiumHD [6]). Segmentation errors further confound single-cell assignments, limiting reconstruction of intercellular signaling.

Single-cell spatial proteomics (e.g., CODEX, MIBI-TOF, IMC, CosMx SMI) offer a complementary view by quantifying surface and signaling proteins, improving segmentation and communication analysis [7, 8, 9, 10]. However, they are limited in characterizing the functional impact of communication and spatial context. Given the feasibility of collecting paired scRNA-seq, integration of scRNA-seq and spatial proteomics benefits from the strengths of each: precise neighborhoods and interactions from proteomics, and fine-grained phenotypes and gene programs downstream of interactions from scRNA-seq. Thus, there is need for robust strategies integrating scRNA-seq and spatial proteomics to expose how local environment and communication shape phenotypes and composition.

Deep generative models for single-cell integration [11, 12, 13, 14, 15] are less suited to spatial proteomics. TotalVI [16] requires paired measurements and barcodes. COVET [17] leverages local covariance, which is less useful for small panels. MIDAS [18] achieves mosaic integration and knowledge transfer, but relies on matching barcodes and does not use spatial information. In spatial protein and transcriptomic integration, “linkage” refers to whether features are measured in or can be predicted by both modalities [19, 20]. Strong-linkage assumptions compress cell heterogeneity and limit transcriptome-wide inference [21, 22, 23, 24, 25]. Weak-linkage approaches like MaxFuse [26] and scMODAL [27] relax feature overlap but rely on gene–protein mappings and ignore spatial context, though such context defines cell states (e.g., T-cell exhaustion in tumor microenvironments [28, 4]).

We present ARCADIA (ARchetype-based Clustering and Alignment with Dual Integrative Autoencoders) which leverages similar cell type/state compositions between modalities and establishes correspondence by matching extreme states inferred as archetypes [28], rather than linked features. ARCADIA is designed for weakly supervised, cross-modal integration settings where harmonized cell types are available, but explicit cell and feature links are absent. ARCADIA trains separate variational autoencoders (VAEs) for scRNA-seq and spatial proteomics and constrains latent spaces so that cells with similar identities co-localize in entangled embeddings. This design does not require barcode pairing, tolerates targeted panels, and explicitly respects spatial organization patterns during integration, while learning phenotypic shifts influenced by spatial context. Dual VAEs support bidirectional knowledge transfer: (i) imputation of spatial niche labels and spatially informed differential expression for scRNA-seq, and (ii) reconstruction of gene programs within and across niches for spatial proteomics, which informs cell communication beyond colocalization.

We demonstrate ARCADIA on a semi-synthetic dataset of paired scRNA-seq and spatial proteomics as well as the only published experimentally paired dataset of scRNA-seq and CODEX from human tonsils in separate studies. In both, ARCADIA outperforms existing methods and recapitulates spatially dependent immune states, linking modalities through composition-level priors without overfitting to sparse panels.

## Materials and Methods

### Overview of ARCADIA

ARCADIA employs a dual variational autoencoder (VAE) framework with coupled objectives that jointly regularize latent representations through shared biological structure inferred from expression patterns of cell types/states across modalities (**Fig. 1A**). Each cell is represented as a convex combination of extreme points, or “archetypes,” and latent distributions of matched archetypes are linked via cross-modal regularization to preserve geometry and encourage biological concordance (**Fig. 1B**). Using archetype-anchored coupling, dual VAEs learn entangled yet modality-specific representations (**Fig. 1C**). ARCADIA’s latent spaces enable bidirectional analysis: how phenotypic variation depends on spatial cell neighborhood and how spatial niches shape transcriptional programs (**Fig. 1D**).

**Figure 1.**
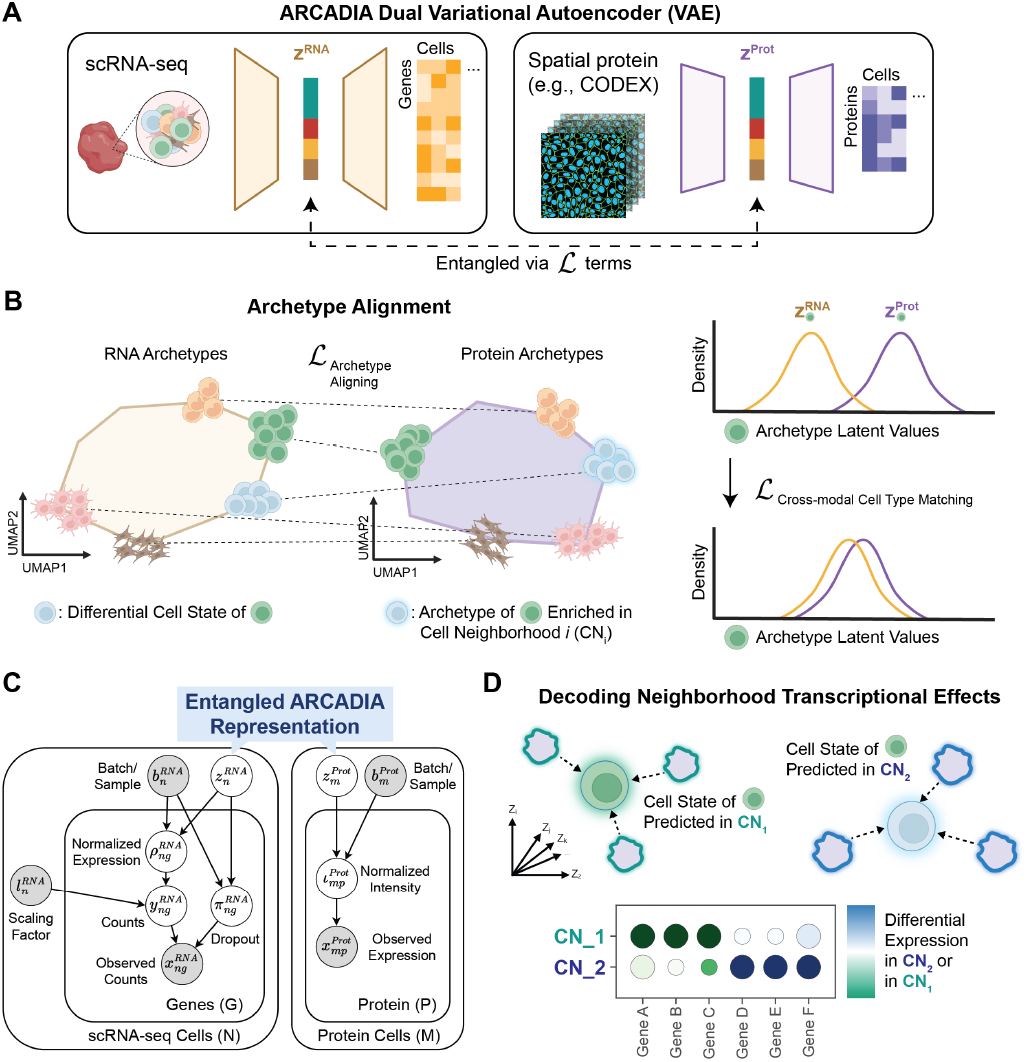
Overview of the ARCADIA Framework. **(A)** Schematic of dual variational autoencoder (VAE) architecture including inputs and outputs. **(B)** ARCADIA aligns archetypes representing cell types/states in both modalities. Archetypes are convex combinations of extreme points on polytopes based on RNA expression (scRNA-seq) or self and neighborhood protein expression (spatial protein). Within matched archetypes, latent distributions are constrained using cross-modal matching loss to yield an entangled representation preserving cell type geometry and concordance across modalities. **(C)** Dual VAE graphical model. Latent variables are circles, and observed variables are shaded circles. **(D)** Data integration and trained dual VAEs allow for investigations into spatial dependencies of cell phenotypes including the effect of CNs on expression. CN: Cell Neighborhood.

### Spatial Protein Features

For each cell, we assemble two feature types: the cell’s own protein-marker expression and spatially-derived aggregate neighborhood features. Neighborhood features are the mean expression of each protein in a spatial neighborhood (using physical coordinates). To ensure locality, long-range edges are pruned by removing connections above the 95th percentile of inter-cell distances, and the 15 nearest neighbors are retained to ensure connectivity. Given the pruned neighbor set 𝒩_*i*_ for cell *i*, the neighborhood-mean feature for protein 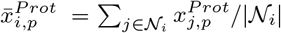 yielding matrix 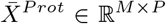 with rows 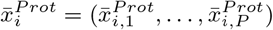.

This matrix is z-standardized (per protein across cells), mitigating heteroscedasticity and improving numerical stability. Neighborhood features are concatenated with protein markers to produce a spatially-aware vector. Spatial features are scaled with an adjustable hyperparameter *α*_*s*_ < 1 prior to concatenation to balance their dynamic range with that of cell-specific expression.

For assessment, cell neighborhood (CN) labels are obtained by K-means clustering (similar to [29]) of neighborhood-mean profiles, providing reference annotations of histological niches. Cluster labels are used exclusively for evaluation of spatial separability, rather than as inputs or supervision for modeling.

### Archetype Analysis and Alignment

ARCADIA introduces an archetype-based alignment framework that establishes biological correspondence between scRNA-seq and spatial proteomics without requiring feature-level linkage. ARCADIA learns modality-specific archetypes that capture *extreme cellular states* and aligns them using *composition-level alignment* derived from cell-type distributions. The specification of cell types for ARCADIA is critical – we discuss the impact and importance of label resolution in **Supplementary Material**.

#### Learning modality-specific archetypes

For each modality *ℓ* ∈ {RNA, Spatial Protein}, input data 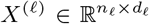 where *n*_*ℓ*_ denotes the number of cells and *d*_*ℓ*_ the number of features, is factorized by Principal Convex Hull Analysis [30] to identify *k* archetypes:

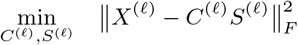

where 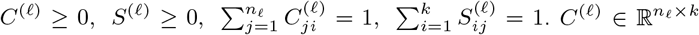 defines archetype-selection weights, and 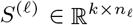 encodes archetype mixture weights for each cell. Each archetype is a convex combination of data points:

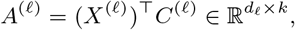

and every cell is represented as a simplex-constrained mixture:

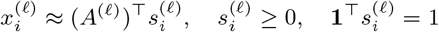

#### Cross-modal archetype alignment

To align archetypes, we compare cell-type composition signatures. For each modality *ℓ*, we compute the archetype-by-cell-type proportion matrix Φ^(*ℓ*)^ ∈ ℝ^*k*×|*T* |^:

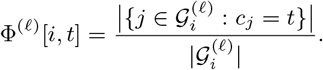

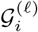 is the set of cells assigned to archetype *i*, based the highest archetype match weight (see **Supplementary Material**), and *c*_*j*_ is the cell-type of cell *j*. We then find the optimal permutation *π*^∗^ that aligns archetypes across modalities by minimizing the cosine distance between their composition profiles:

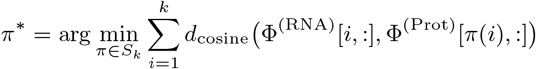

where *S*_*k*_ is the set of all permutations of *k* elements. We solve this optimization using the Hungarian method in 𝒪 (*k*^3^) time [31]. The optimal *k*^∗^ minimizes the average alignment cost: 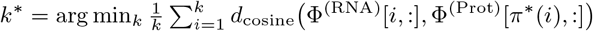.

#### Extreme cell-state anchors

Cells whose archetype mixture vectors 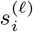 are dominated by a single archetype, 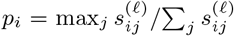, are denoted *anchors* and used as high-confidence archetypes for modality alignment. Anchors represent biologically distinct cell states enabling robust archetype-level alignment prior to latent-space training.

In contrast to feature-level alignment methods, ARCADIA defines cross-modal correspondence via archetype matching, establishing interpretable links between transcriptional and spatial cell states as the biologically grounded foundation for dual-VAE training. Further details on archetype analysis and cross-modal alignment are in the **Supplementary Material**.

### Model Architecture and Training

Both the RNA and protein encoders project to respective 60-dimensional latent spaces. This larger dimensionality compared to [26, 27] reflects high-dimensional inputs: highly variable genes without filtering to protein markers for RNA, and markers concatenated with neighborhood features for protein. Both encoders employ batch normalization and dropout (*ρ* = 0.1), and decoders mirror each encoder. A single AdamW optimizer handles trainable parameters, updating both VAEs in each pass.

For RNA we use a zero-inflated negative binomial (ZINB) likelihood on counts. For protein we use a Gaussian likelihood. We select likelihoods by fitting candidates to random samples from each modality and evaluating Akaike Information Criterion (AIC). Batch labels condition decoders during training, allowing VAEs to absorb batch effects in the latent space. Combined with per-batch archetype alignment, this yields consistent representations across batches and modalities.

As VAEs encode/decode independently, trained VAEs support cross-modal inference (similar to generating counter-factuals in [32]). Additional model details (e.g., generative and inference processes) are in **Supplementary Material**.

### Objective Functions

Dual VAEs learn latent spaces by optimizing four objectives. Modality-specific evidence lower bounds (ELBOs) ensure faithful feature reconstruction while regularizing latent geometry. Structure-preservation loss maintains cell-type relationships and organization so latent geometry reflects known biological structure in both modalities. Cross-modal Maximum Mean Discrepancy (MMD) loss encourages mixing of cell types, promoting correspondence without feature-level alignment. Anchor-guided loss preserves intra-cell-type variation, preventing over-mixing and ensuring phenotypic diversity within cell types is retained. Together, the loss terms entangle latent spaces while enabling cross-modal inference.

#### Evidence lower bound (ELBO)

For modality *ℓ* ∈ {RNA, Spatial Protein}, we define the ELBO:

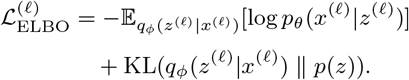

*x*^(*ℓ*)^ is observed data, *z*^(*ℓ*)^ is the latent representation, *q*_*ϕ*_ is the variational encoder, and *p*_*θ*_ is the decoder. We employ standard Kullback–Leibler divergence with normal priors *p*(*z*) = 𝒩 (0, *I*).

#### Anchor-guided cross-modal matching

*Anchors* are cells whose archetype vectors 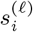 are dominated by a single archetype, so they serve as high-confidence alignment references. The matching loss penalizes distances between anchors via softplus with adaptive margins:

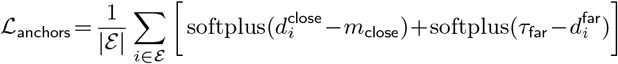

where ℰ is the set of anchors in modality 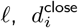 and 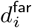 are latent-space distances from anchor *i* to its most and least similar counterpart in the other modality, *m*_close_ = *α*_close_*σ* is a closeness margin (0 < *α*_close_ ≪ 1), and *τ*_far_ = *α*_far_*σ* is a farness margin (*α*_far_ *>* 1), with *σ* as the standard deviation of latent distances. 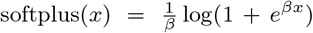 provides smooth, non-negative penalties. Critically, when distances fall within acceptable thresholds (i.e., close enough or far enough), the penalty is near zero, so well-matched pairs to contribute minimally while poorly-matched pairs incur larger penalties. This design encourages natural alignment without hard constraints.

#### Cell-type structure preservation

For each modality, cell-type centroids are computed in archetype space and VAE latent space. Structure preservation loss aligns inter-cell-type affinity matrices computed via Gaussian kernels:

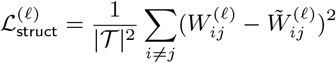

where 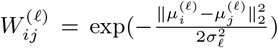 is the affinity between cell types *i* and *j* in latent space with centroids 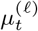, and 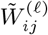 is the corresponding affinity in archetype space with centroids 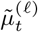. Additional cohesion and separation losses promote cluster compactness and inter-cluster distinctness.

#### Cross-modal alignment

We compute the Maximum Mean Discrepancy (MMD) between distributions of cell types in each latent *z*^(*ℓ*)^ to align them. Minimizing MMD ensures cell types occupy similar regions in *z*^(*ℓ*)^ while facilitating mixing and preventing modality-specific artifacts. For each cell type *t* ∈ 𝒯, 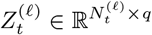 denotes the matrix of latent representations in modality *ℓ*, where 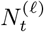 is the number of type-*t* cells and *q* is the latent dimension. Each row of 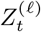 is a cell’s latent embedding. The MMD is:

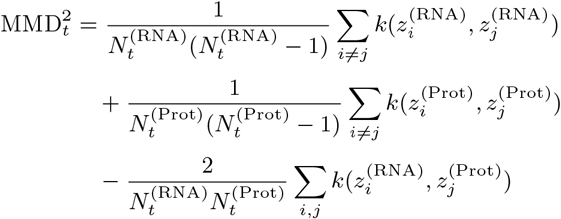

where the RBF kernel is 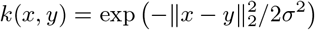 and *σ* is adaptively set via the median heuristic on combined data. Specifically, 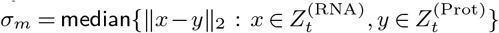 scales the kernel automatically to the data’s natural distance distribution. Instead of a single bandwidth, an ensemble of MMDs [33] is computed at multiple scales: fine- 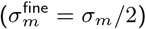, medium- 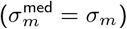, and coarse-grained 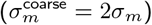:

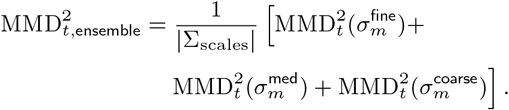

Fine-grained scales detect tightly clustered cell types, medium scales capture intermediate structure, and coarse-grained scales capture loosely distributed ones, ensuring alignment respects differences in how cell types disperse in the latent space. Cross-modal alignment loss is the average MMD across cell types:

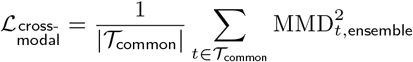

#### Total training objective

The total loss combines components via fixed hyper-parameters:

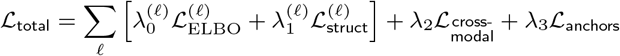

where 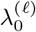 and 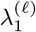 control reconstruction and structure preservation and *λ*_2_ and *λ*_3_ control alignment strength. Further details on objective functions are in **Supplementary Material**.

## Experiments and Results

### Datasets

#### Semi-Synthetic Dataset

To define ground-truth spatial niches dictating cell states, we took an expertly annotated CITE-seq dataset with paired protein and RNA expression [34], split it into two disjoint sets of protein- and RNA-expressing cells, and positioned protein cells on a spatial grid where each quadrant contained certain subtypes. Specifically, we placed different B cells subtypes in cell neighborhoods (CNs), surrounded either by no other cell types or by T and dendritic cells. We held out spatial context information for RNA cells. Protein profiles alone could not describe the range of B cell states, but could be linked to RNA profiles to describe these states. As such, we tested whether ARCADIA learns cross-modal cell types based on aligned archetypes as well as spatially-dependent variation (**Fig. 2A**).

**Figure 2.**
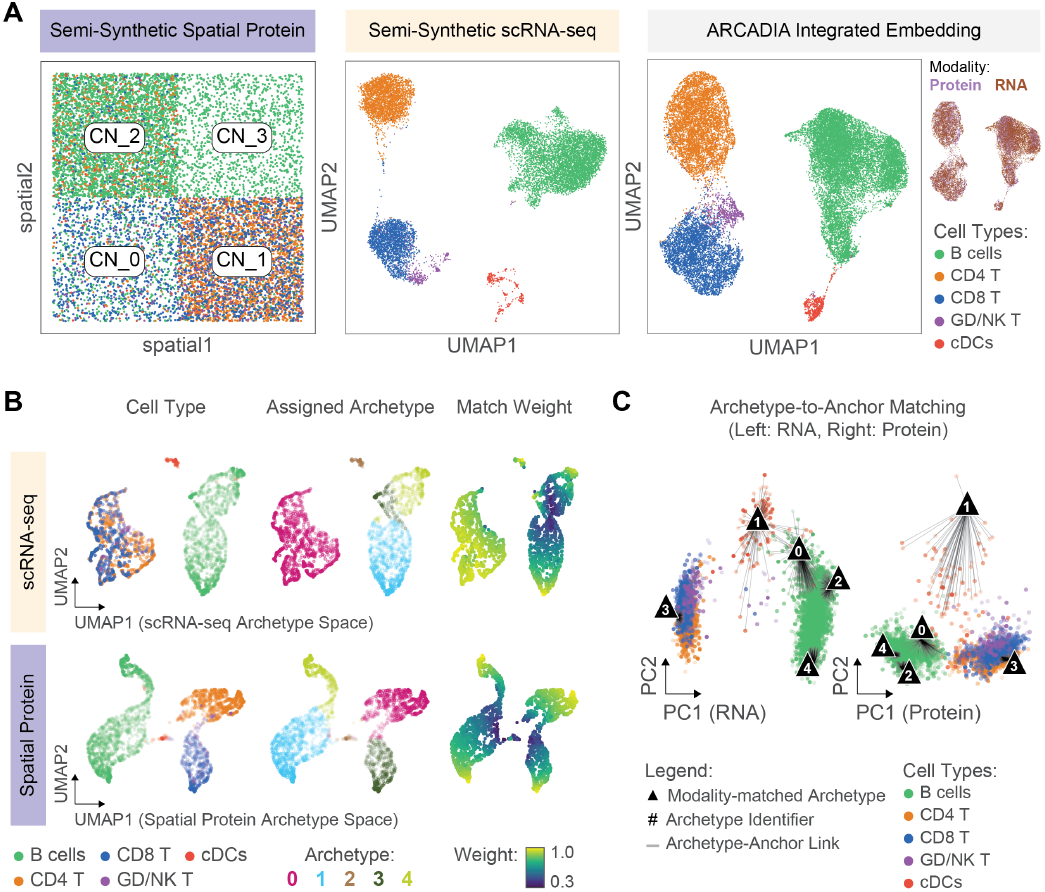
Semi-synthetic data integration and archetype alignment. **(A)** Left, spatial grid of protein cells and UMAP of RNA cells colored by type; right, ARCADIA embedding colored by type (outer) and modality (inner). **(B)** RNA and protein cells in archetype space. Opacity reflects match weight. **(C)** Archetype matching for RNA and protein cells in PC space. CN: Cell Neighborhood.

#### Human Tonsil Dataset

To demonstrate the utility of ARCADIA to discover spatially-dependent cell state patterns, we integrated human tonsil data collected in separate studies [35, 36, 37]. At the time of this study, this collection is the only publicly available, experimentally-paired scRNA-seq and CODEX dataset. Both modalities were already expertly-annotated. As in [29], we characterized 10 cell neighborhoods (CNs) based on protein expression of each cell and its surrounding neighbors.

#### Data Preprocessing

We performed standard scRNA-seq and protein data quality control, feature selection, and normalization. We also harmonized cell type labels, which is crucial for archetype alignment. Preprocessing protocols, including dataset-specific considerations, are in **Supplementary Material**.

### Computational Benchmarking

We assessed ARCADIA’s scalability using the semi-synthetic CITE-seq dataset (∼30k total cells) and the human tonsil dataset (∼180k total cells). Preprocessing runtime, including spatial graph construction and archetype generation, scaled with dataset size. In contrast, dual-VAE training (300 epochs) was comparable (1.3 vs. 1.8 hours), as cost is driven by model complexity. Crucially, peak GPU memory usage was low for both datasets (3.0–3.2 GB), demonstrating ARCADIA can process large spatial datasets on standard hardware (NVIDIA T4) (**Table 2**). Detailed hardware specifications, complexity analysis, and resource profiling are in the **Supplementary Material**.

**Table 1.**
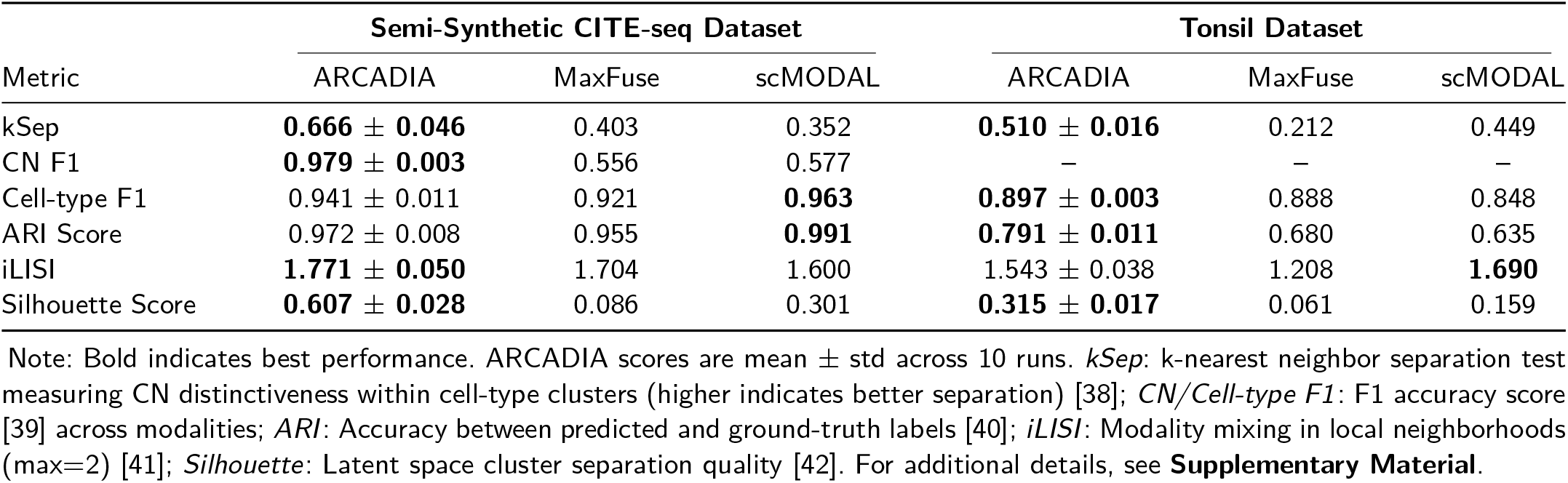
Dataset Integration Performance Benchmarking.

| Metric | Semi-Synthetic CITE-seq Dataset |  |  | Tonsil Dataset |  |  |
| --- | --- | --- | --- | --- | --- | --- |
|  | ARCADIA | MaxFuse | scMODAL | ARCADIA | MaxFuse | scMODAL |
| kSep | **0.666 ± 0.046** | 0.403 | 0.352 | **0.510 ± 0.016** | 0.212 | 0.449 |
| CN F1 | **0.979 ± 0.003** | 0.556 | 0.577 | — | — | — |
| Cell-type F1 | 0.941 ± 0.011 | 0.921 | **0.963** | **0.897 ± 0.003** | 0.888 | 0.848 |
| ARI Score | 0.972 ± 0.008 | 0.955 | **0.991** | **0.791 ± 0.011** | 0.680 | 0.635 |
| iLISI | **1.771 ± 0.050** | 1.704 | 1.600 | 1.543 ± 0.038 | 1.208 | **1.690** |
| Silhouette Score | **0.607 ± 0.028** | 0.086 | 0.301 | **0.315 ± 0.017** | 0.061 | 0.159 |

Note: Bold indicates best performance. ARCADIA scores are mean ± std across 10 runs. *kSep*: k-nearest neighbor separation test measuring CN distinctiveness within cell-type clusters (higher indicates better separation) [38]; *CN/Cell-type F1*: F1 accuracy score [39] across modalities; *ARI*: Accuracy between predicted and ground-truth labels [40]; *iLISI*: Modality mixing in local neighborhoods (max=2) [41]; *Silhouette*: Latent space cluster separation quality [42]. For additional details, see **Supplementary Material**.

**Table 2.** Dataset Statistics and Computational Benchmarks.

| Metric | CITE-seq | Tonsil |
| --- | --- | --- |
| **Dimensions (cells × genes or features)** |  |  |
| scRNA-seq | 15,337 × 1,796 | 12,917 × 2,000 |
| Spatial Protein | 14,021 × 220 | 169,468 × 92 |
| **Preprocessing (CPU)** |  |  |
| Runtime | 13.5 min | 56.9 min |
| Peak System RAM | 14.9 GB | 38.6 GB |
| **Training (GPU, 300 Epochs)** |  |  |
| Trainable Parameters | ~15.3 Million | ~16.0 Million |
| Runtime | 1.3 h | 1.8 h |
| Peak System RAM | 7.7 GB | 41.7 GB |
| Peak GPU VRAM | 3.0 GB | 3.2 GB |
| Avg. CPU Usage | 197% (~2 cores) | 190% (~2 cores) |

Note: Benchmarks using 8-core Intel Xeon CPU and 15 GB VRAM NVIDIA Tesla T4 GPU. For additional details, see **Supplementary Material**.

### Performance on Semi-Synthetic Dataset

ARCADIA exhibited robust and meaningful archetype matching in both the RNA and protein expression space (**Fig. 2B,C**) and clear separations by cell type with even distribution of cells across modalities in the integrated embedding space (**Fig. 2A, Fig. S1A**). Crucially, beyond co-embedding immune populations, ARCADIA captured the spatial patterning of B-cell subtypes across CNs (**Fig. S2A,B**), evidenced by clear cell clusters from different CNs in the integrated embedding (**Fig. S2C**). MaxFuse [26] and scMODAL [27] integrated cell types, but could not learn spatially-dependent cell state patterns (**Fig. S2C**).

ARCADIA out-performed both MaxFuse and scMODAL in most benchmarking metrics, while all three comparably predicted cell types (**Table 1, Fig. 3A**). For each method we used integrated embeddings to predict CNs from scRNA-seq and compared them to ground-truth, to test how each method could recover cell subtypes from spatial grid location (in protein) and phenotype (in RNA). ARCADIA significantly outperformed both MaxFuse and scMODAL using CN F1 scores (**Table 1**). In particular, ARCADIA accurately predicted spatial location for B cell subtypes in CN 1 and CN 2, which had the greatest density and subtype diversity (**Fig. 3B, Fig. S2A,B**). See **Table S1** in **Supplementary Material** for additional benchmarking.

**Figure 3.**
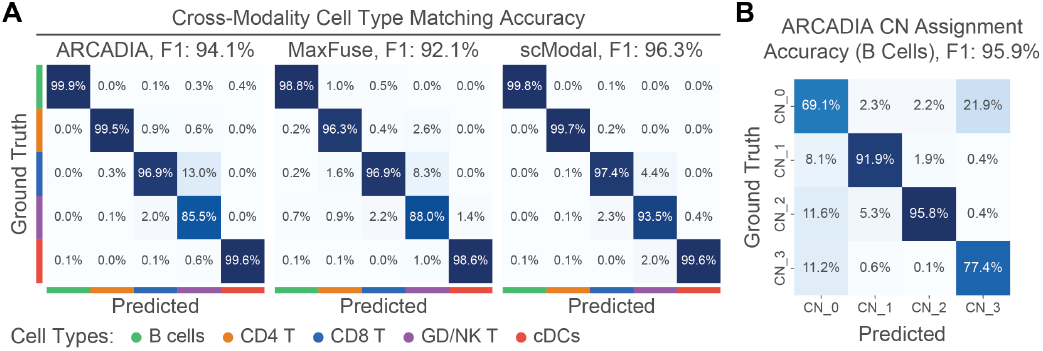
Integration performance on semi-synthetic data. **(A)** Cell type confusion matrices for ARCADIA, MaxFuse, and scMODAL. **(B)** Confusion matrices for B cell-specific CN prediction from ARCADIA.

### Performance on Human Tonsil Dataset

Next we studied spatially-dependent behavior of immune cells in human tonsil samples. ARCADIA recovered both major immune-cell populations (**Fig. 4A**), and cell-state gradients across spatial contexts (**Fig. 4B, Fig. S3A**). While MaxFuse and scMODAL performed comparably in broad cell type integration (**Fig. S3B, Table 1**), only ARCADIA captured spatially-dependent phenotypic variation in B, T, and dendritic cells (**Fig. S3C**).

**Figure 4.**
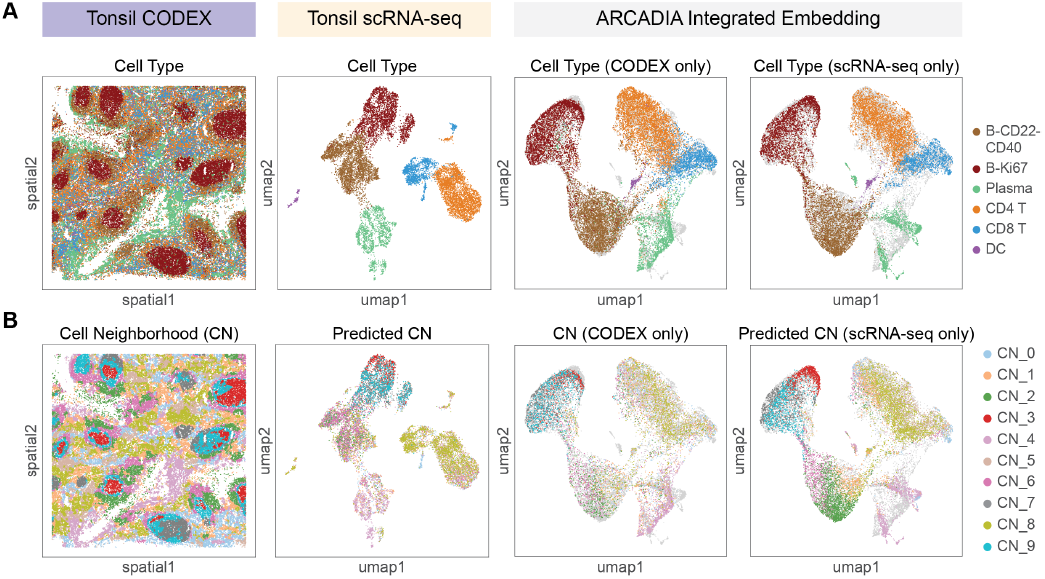
ARCADIA integrates human tonsil scRNA-seq and CODEX data. **(A)** Cell types in the CODEX tonsil tissue sample (left), in scRNA-seq (middle), and in ARCADIA space (right). **(B)** Computed cell neighborhoods (CNs) in the CODEX tissue sample (left), predicted CN from scRNA-seq (middle), and ground-truth and predicted CN in ARCADIA space (right). DC: Dendritic cell.

Previous studies characterized spatial structures of germinal-centers and peripheral regions in the tonsil [29]. ARCADIA’s integrated embedding links these spatial (sub)structures (represented by CNs) to the phenotypic spectrum of states of B, T, and dendritic cells. In predicting CN for each cell, ARCADIA allows for spatially-informed transcriptomic analysis at the level of cells within CNs, as opposed to the level of CNs as in [29]. We next sought to use the ARCADIA framework to contrast states of B, CD8 T, and CD4 T cells within different spatial regions.

### ARCADIA Learns Spatially-Resolved Gene Expression Patterns

We used ARCADIA to predict CN membership for each scRNA-seq cell. Subsequently, we performed differentially expressed gene (DEG) analysis within cell types but across inferred CNs, thus unbiasedly studying how, within cell populations, phenotypic states vary by CN in the tonsil. DEG testing was restricted to CNs with adequate cell type representation (**Fig. S3A**). Complete lists of top DEGs are in **Figs. S4, S5, and S6**.

B cells in central germinal-center (GC) CN 3 expressed hypermutation (*BCL6, AICDA, MYBL1, BACH2*), activation/BCR signaling (*CD84, MME, LRMP, CXCR4*), and proliferation markers (*MKI67, BIRC5*). Peripheral CN 7 and CN 9 B cells expressed plasma-cell-like (*CD83, FCRL5*), metabolic (*GAPDH, LDHA, ODC1*), and antigen-presentation (*HLA-A, HLA-DQA1, HLA-DQB1*) genes (**Fig. 5A**). These results suggest B cells deep within GCs proliferate/hypermutate, whereas in outer GC zones they undergo terminal differentiation and immune-surveillance.

**Figure 5.**
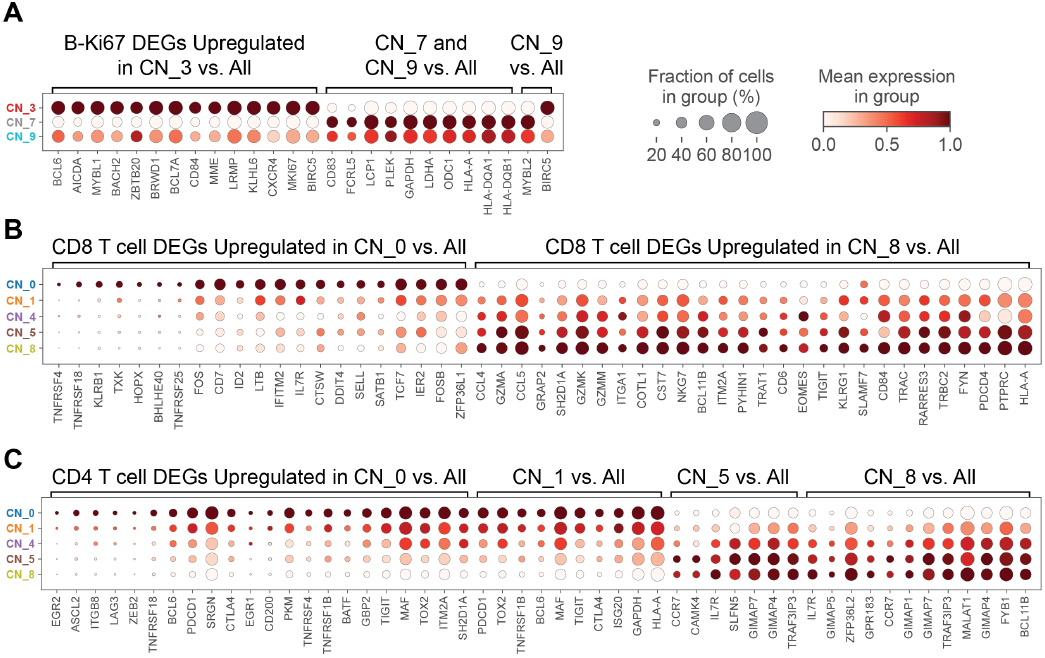
Spatially-resolved differential expression. Genes expressed by B **(A)**, CD8 T **(B)**, and CD4 T cells **(C)** predicted to originate from specific CNs. Only CNs with sufficient representation are shown (see Fig. S3A). Expression is scaled across CNs. Full DEGs in Figs. S4–S6.

CD8 T cells in CN 0 had effector (*TNFRSF4, CTSW*), activated (FOS, LTB), and stem-like (*IL7R, TCF7*) states, while those in CN 8 co-expressed cytotoxic (*CCL4, GZMA*) and exhaustion (*TIGIT, KLRG1, PDCD4*) markers (**Fig. 5B**). CD4 T cells in CN 4 showed terminally exhausted phenotypes (*ZEB2, LAG3, CTLA4*), whereas CN 1 displayed T follicular helper (Tfh)-like states (*BCL6, MAF, GAPDH, HLA-A*) and CN 5/8 contained stem-like precursors (*IL7R, CCR7*) (**Fig. 5C**). Additional DEGs and biological implications are in **Supplementary Material**.

Thus, ARCADIA reveals coordinated B-cell maturation and T-cell activation/exhaustion dependent on spatial niche relative to GCs, recapitulating canonical B and Tfh cell biology [43, 44, 45, 46] and understudied follicular CD8 T-cell behaviors [47].

## Discussion and Future Work

As additional spatial proteomics data become available with paired scRNA-seq, ARCADIA provides a unified framework to probe relationships between tissue microenvironment and cell state, including in cases where ST fails (e.g., limited panels, poor segmentation). Benchmarks on multiple datasets show that ARCADIA outperforms weak-linkage methods and uniquely recovers spatially-dependent functional states across immune lineages. We note model accuracy depends on balanced cell-type representation across data, since archetype alignment relies on shared composition. As with other deep generative models, careful hyperparameter tuning is required to maintain biologically meaningful separation of subtle states. Future work will extend cross-modal decoding (e.g., predicting RNA expression from CODEX). In sum, ARCADIA establishes a generative, interpretable bridge between scRNA-seq and spatial proteomics and bidirectional modeling of interactions and tissue remodeling.

## Acknowledgments

We thank Mingxuan Zhang, Joshua Meyers, Yinuo Jin, Ady Zhang, José McFaline-Figueroa, and Andrew J. Blumberg for helpful discussions and feedback.

## Supplementary Data

Supplementary data and figures are included in the submission.

## Competing Interests

No competing interests are declared.

## Funding

K.H.H. was supported by the NSF GRFP under Grant No. DGE-2036197. E.A. is supported by the NIH NHGRI grant R01HG012875, R21HG012639, and grant number 2022-253560 from the Chan Zuckerberg Initiative DAF, an advised fund of Silicon Valley Community Foundation.

## Data Availability

All data analyzed are previously published. Source code is accessible at https://github.com/azizilab/ARCADIA_public. Reproducibility scripts and data are available at https://github.com/azizilab/arcadiareproducibility.

## Notes

### Competing Interest Statement

The authors have declared no competing interest.

### Summary of Updates

Updated manuscript with slight language edits

## References

1. Patrik L. Ståhl et al. Visualization and analysis of gene expression in tissue sections by spatial transcriptomics. Science, 2016.

2. E. Armingol et al. Deciphering cell–cell interactions and communication from gene expression. Nat Rev Genet, 2021.

3. E. Lundberg et al. Spatial proteomics: a powerful discovery tool for cell biology. Nat Rev Mol Cell Biol, 2019.

4. Itay Tirosh et al. Dissecting the multicellular ecosystem of metastatic melanoma by single-cell rna-seq. Science, 2016.

5. E. Azizi et al. Single-cell map of diverse immune phenotypes in the breast tumor microenvironment. Cell, 2018.

6. Lingting Shi et al. Spatiotemporal single-cell analysis reveals t cell clonal dynamics& bioRxiv, 2025.

7. Yury Goltsev et al. Deep profiling of mouse splenic architecture with codex multiplexed imaging. Cell, 2018.

8. L. Keren et al. Mibi-tof: A multiplexed imaging platform relates cellular phenotypes and tissue structure. Sci Adv, 2019.

9. C. Giesen et al. Highly multiplexed imaging of tumor tissues with subcellular resolution by mass cytometry. Nat Methods, 2014.

10. S. He et al. High-plex imaging of rna and proteins at subcellular resolution in fixed tissue by spatial molecular imaging. Nat Biotechnol, 2022.

11. Romain Lopez et al. Deep generative modeling for single-cell transcriptomics. Nature Methods, 2018.

12. K.E. Wu et al. Babel enables cross-modality translation between multiomic profiles at single-cell resolution. Proc. Natl. Acad. Sci. U.S.A., 2021.

13. J. Zhao et al. Adversarial domain translation networks for integrating large-scale atlas-level single-cell datasets. Nat Comput Sci, 2022.

14. ZJ. Cao et al. Multi-omics single-cell data integration and regulatory inference with graph-linked embedding. Nat Biotechnol, 2022.

15. Z. Tang et al. Modal-nexus auto-encoder for multi-modality cellular data integration and imputation. Nat Commun, 2024.

16. A. Gayoso et al. Joint probabilistic modeling of single-cell multi-omic data with totalvi. Nat Methods, 2021.

17. D. Haviv et al. The covariance environment defines cellular niches for spatial inference. Nat Biotechnol, 2025.

18. Z. He et al. Mosaic integration and knowledge transfer of single-cell multimodal data with midas. Nat Biotechnol, 2024.

19. R. Argelaguet et al. Computational principles and challenges in single-cell data integration. Nat Biotechnol, 2021.

20. Y. Xu et al. Diagonal integration of multimodal single-cell data: potential pitfalls and paths forward. Nat Commun, 2022.

21. Joshua D. Welch et al. Single-cell multi-omic integration compares and contrasts features of brain cell identity. Cell, 2019.

22. Tim Stuart et al. Comprehensive integration of single-cell data. Cell, 2019.

23. Jinzhuang Dou et al. Unbiased integration of single cell multi-omics data. bioRxiv, 2020.

24. J. Dou et al. Bi-order multimodal integration of single-cell data. Genome Biol, 2022.

25. B. Zhu et al. Robust single-cell matching and multimodal analysis using shared and distinct features. Nat Methods, 2023.

26. S. Chen et al. Integration of spatial and single-cell data across modalities with weakly linked features. Nat Biotechnol, 2024.

27. G. Wang et al. scmodal: a general deep learning framework for comprehensive single-cell multi-omics data alignment with feature links. Nat Commun, 2025.

28. S. He et al. Starfysh integrates spatial transcriptomic and histologic data to reveal heterogeneous tumor–immune hubs. Nat Biotechnol, 2025.

29. Daisy Y. Ding et al. Quantitative characterization of tissue states using multiomics and ecological spatial analysis. Nature Genetics, 2025.

30. Adele Cutler et al. Archetypal analysis. Technometrics, 1994.

31. Harold W. Kuhn. The hungarian method for the assignment problem. Naval Research Logistics Quarterly, 1955.

32. P. Boyeau et al. Deep generative modeling of sample-level heterogeneity in single-cell genomics. Nat Methods, 2025.

33. Antonin Schrab et al. Mmd aggregated two-sample test. Journal of Machine Learning Research, 2023.

34. Yuhan Hao et al. Integrated analysis of multimodal single-cell data. Cell, 2021.

35. J. Kennedy-Darling et al. Highly multiplexed tissue imaging using repeated oligonucleotide exchange reaction. Eur. J. Immunol., 2021.

36. Hamish W. King et al. Single-cell analysis of human b cell maturation predicts how antibody class switching shapes selection dynamics. Sci Immunol, 2021.

37. Hamish W. King et al. Integrated single-cell transcriptomics and epigenomics reveals strong germinal center–associated etiology of autoimmune risk loci. Sci Immunol, 2021.

38. Maren Büttner et al. A test metric for assessing single-cell rna-seq batch correction. Nature Methods, 2019.

39. Cornelis Joost van Rijsbergen. Information Retrieval. Butterworths, 1979.

40. Lawrence Hubert et al. Comparing partitions. Journal of Classification, 1985.

41. Ilya Korsunsky et al. Fast, sensitive and accurate integration of single-cell data with harmony. Nature Methods, 2019.

42. Peter J. Rousseeuw. Silhouettes: A graphical aid to the interpretation and validation of cluster analysis. Journal of Computational and Applied Mathematics, 1987.

43. Domenick E. Kennedy et al. Compartments and connections within the germinal center. Frontiers in Immunology, 2021.

44. U. Klein et al. Germinal centres: role in b-cell physiology and malignancy. Nat Rev Immunol, 2008.

45. N. De Silva et al. Dynamics of b cells in germinal centres. Nat Rev Immunol, 2015.

46. S. Crotty. Follicular helper cd4 t cells (tfh). Annu Rev Immunol, 2011.

47. CH. Koh et al. Cd8 t-cell subsets: heterogeneity, functions, and therapeutic potential. Exp Mol Med, 2023.

