## Supplementary Material for "ARCADIA Reveals Spatially Dependent Transcriptional Programs through Integration of scRNA-seq and Spatial Proteomics"

#### S1 Detailed Data Preprocessing

ARCADIA requires careful preprocessing of both scRNA-seq and spatial proteomics data to ensure biological signal preservation while removing technical artifacts. The preprocessing pipeline harmonizes cell-type annotations, filters low-quality measurements, normalizes expression values, and constructs spatially-aware feature representations that capture both cell-intrinsic and neighborhood-level information. In this work, we apply this preprocessing methodology to two distinct multimodal datasets: Semi-synthetic CITE-seq data [1] consisting of transcriptomics with paired protein barcodes and synthetically added spatial information, and human tonsil data [2, 3, 4] consisting of independently collected scRNA-seq and spatial proteomics. Dataset-specific adaptations are described separately below.

##### S1.1 Quality Control and Cell Filtering

**RNA quality control.** For scRNA-seq data, we apply standard quality control metrics to remove low-quality cells and noisy genes. Specifically, we filter cells with fewer than 200 detected genes (genes with non-zero counts) and genes expressed in fewer than 3 cells. Additionally, for each cell we compute total UMIs expressed and mitochondrial gene expression percentage, and we filter cells with fewer than 1,000 UMIs expressed and mitochondrial gene expression percentage above 20% to remove low-quality cells with likely RNA degradation or excessive mitochondrial contamination.

**Protein quality control.** For spatial proteomics data, we apply minimum detection thresholds to ensure data quality: cells must express at least $p_{\min} = 25$ markers above zero.

**Median absolute deviation outlier removal for protein data.** To remove outlier cells within each cell type while preserving biological heterogeneity in the protein modality, we employ cell-type-aware outlier detection using the median absolute deviation (MAD) criterion. For each cell type $t \in \mathcal{T}^{(\text{Prot})}$, let $S_t = \{i : c_i = t\}$ denote the set of protein cells assigned to type $t$, where $c_i$ is the cell-type label for cell $i$. For protein marker intensity sums or quality metrics, we compute the median value:

$$\tilde{m}_t = \text{median}\{m_i : i \in S_t\},$$

where $m_i$ represents a cell-level quality metric (e.g., total marker intensity). The MAD is:

$$\text{MAD}_t = \text{median}\left\{ \left| m_i - \tilde{m}_t \right| : i \in S_t \right\}.$$

A protein cell $i \in S_t$ is flagged as an outlier if

$$m_i < \tilde{m}_t - \tau_{\text{MAD}} \cdot \text{MAD}_t \quad \text{or} \quad m_i > \tilde{m}_t + \tau_{\text{MAD}} \cdot \text{MAD}_t,$$

where $\tau_{\text{MAD}} = 2$ is the threshold parameter. This method removes cell-type outliers by using each cell type's own population statistics as a more faithful representation of its distribution, rather than relying on a global average across all cells.

#### S1.2 Cell Type Harmonization and Filtering

**Cross-modal cell-type alignment.** To ensure accurate archetype matching across modalities, we harmonize cell-type annotations by mapping modality-specific labels to a common set of categories. After mapping, we retain only cells that belong to cell types present in both modalities, ensuring consistent cell-type definitions across the RNA-seq and spatial proteomics data for downstream integration analysis.

#### S1.3 RNA Normalization

**Library size normalization and log transformation.** We scale to a target library size of 40,000 for each cell to account for varying sequencing depths. We also apply a log-plus-one transformation to the normalized expression. We use log-normalized expression for visualization, principle component computation, and differential expression; for VAE training, we retain count data as input, as the zero-inflated negative binomial likelihood explicitly models whole number counts.

#### S1.4 Gene Selection and Dimensionality Reduction

**Highly variable gene selection.** We select highly variable genes (HVGs) using a variance-stabilized procedure applied to raw counts with batch awareness, i.e., `scanpy.pp.highly_variable_genes` with `flavor='seurat_v3'`. We initially select the top 2,000 genes ranked by normalized dispersion. To determine the optimal number of features, we further apply the Kneedle algorithm [5] to the log-transformed variance curve of the ranked HVGs. Given the sorted variances $\sigma_1^2 \geq \sigma_2^2 \geq \cdots \geq \sigma_{2000}^2$, Kneedle identifies the index $n_{\text{knee}}$ where the curve transitions from steep to flat by detecting maximum curvature in the normalized curve $y_i = \log(\sigma_i^2)$. The final HVG set size is thus $n_{\text{HVG}} = \min(2000, n_{\text{knee}})$.

**Principal component analysis.** During archetype discovery, we project both modalities into lower-dimensional subspaces via principal component analysis (PCA). For RNA data, we compute PCs on the HVG-filtered, log-normalized expression matrix, retaining $K^{(\mathrm{RNA})} = 50$ PCs that capture the dominant axes of cell type/state variation. For proteomic data, after concatenating normalized intrinsic and neighborhood protein features we compute and retain $K^{(\mathrm{Prot})} = 30$ PCs.

##### S1.5 Cross-Modal Dataset Balancing

**Proportion-aware subsampling.** ARCADIA archetype matching relies on cell-type proportions matching across modalities, so we apply proportion-aware subsampling to ensure consistent cell-type distributions between RNA and spatial proteomics datasets. Let $n_t^{(\mathrm{RNA})}$ and $n_t^{(\mathrm{Prot})}$ denote the counts of cell type $t$ in the respective modalities. We compute cell-type proportions in the smaller dataset (by total cell count) as

$$p_t^{(\mathrm{smaller})} = \frac{n_t^{(\mathrm{smaller})}}{\sum_{t'} n_{t'}^{(\mathrm{smaller})}}.$$

To determine the maximum balanced dataset size, we identify the limiting cell type $t^*$ in the larger dataset:

$$t^* = \underset{t}{\mathrm{argmin}}\, \frac{n_t^{(\mathrm{larger})}}{p_t^{(\mathrm{smaller})}},$$

where $n_t^{(\mathrm{larger})}$ is the count of type $t$ in the larger modality. The maximum balanced size is then

$$N_{\max} = \frac{n_{t^*}^{(\mathrm{larger})}}{p_{t^*}^{(\mathrm{smaller})}}.$$

For each cell type $t$, we randomly subsample the larger dataset (without replacement) to

$$n_t^{(\mathrm{target})} = \min\big(\lfloor p_t^{(\mathrm{smaller})} \cdot N_{\max} \rfloor,\ n_t^{(\mathrm{larger})}\big).$$

We perform this subsampling independently for each modality so that both modalities are reduced to common cell-type distributions, thus ensuring archetype correspondence and preventing artificial matches driven by distributional imbalance rather than biological similarity.

#### S2 Detailed Archetypal Analysis

The goal of this archetype analysis is to identify corresponding cells between modalities by projecting cell from both modalities to a unified archetype coordinate system. The overview of steps is as follows:

1. Generate archetypes separately for each modality (and per batch if present) to obtain modality-specific vector bases.

2. Align archetypes across modalities by comparing corresponding cell-type proportions.

3. Represent every cell as a linear combination of archetypes, thus creating a unified archetype embedding across modalities.

4. Select cells whose archetype coefficients are dominated by a single archetype as *anchors*, representing high-confidence cells the will guide the cross-modality alignment.

5. Compute cross-modal cell–cell distances in archetype space to measure similarity between RNA and protein cells.

##### Step 1: Construct archetypes

We build modality-specific archetypes per batch using Principal Convex Hull Analysis (PCHA) [6] on low-dimensional embeddings (e.g., PCA) to reduce complexity commonly found in spatial genomics data [7].

**Archetypal analysis formulation.** Let $X^{(\mathrm{RNA})} \in \mathbb{R}^{N\times G}$ denote the RNA expression matrix with $N$ cells and $G$ genes, and let $X^{(\mathrm{Prot})} \in \mathbb{R}^{M\times P}$ denote the spatial proteomics matrix with $M$ cells and $P$ protein markers. For each modality $\ell \in \{\mathrm{RNA}, \mathrm{Spatial\ Protein}\}$, define $C^{(\ell)} \in \mathbb{R}^{n_\ell\times k}$ as the archetype-selection matrix, and $S^{(\ell)} \in \mathbb{R}^{k\times n_\ell}$ as the reconstruction-weight matrix, where $n_\ell \in \{N, M\}$ is the number of cells in modality $\ell$ and $k$ is the number of archetypes.

For each modality, we fit archetypes by minimizing the Frobenius-norm reconstruction error:

$$\begin{aligned}
\min_{C^{(\ell)},S^{(\ell)}} \quad & \left\| X^{(\ell)} - C^{(\ell)} S^{(\ell)} \right\|_F^2 \\
\text{subject to} \quad & C^{(\ell)} \geq 0, \quad \sum_{j=1}^{n_\ell} C_{ji}^{(\ell)} = 1 \quad \text{for } i = 1,\dots,k, \\
& S^{(\ell)} \geq 0, \quad \sum_{i=1}^{k} S_{ij}^{(\ell)} = 1 \quad \text{for } j = 1,\dots,n_\ell.
\end{aligned}$$

The archetype matrix for modality $\ell$ is given by

$$A^{(\ell)} = \left(X^{(\ell)}\right)^\top C^{(\ell)} \in \mathbb{R}^{d_\ell\times k},$$

where $d_\ell \in \{G, P\}$, so that each column of $A^{(\ell)}$ is a convex combination of modality-specific cells. Each cell $j$ in modality $\ell$ is then represented as a convex combination of archetypes,

$$X_{j\cdot}^{(\ell)} \approx \left(A^{(\ell)}\right)^\top s_j^{(\ell)},$$

where $s_j^{(\ell)} \in \mathbb{R}^k$ denotes the $j$-th column of $S^{(\ell)}$ and encodes the archetype weight vector for that cell. Together, $C^{(\ell)}S^{(\ell)}$ reconstructs the original modality-specific data matrix for scRNA-seq counts or spatial proteomics features.

##### Step 2: Select optimal cross-modal archetype set

Because the optimal number of archetypes is unknown *a priori*, we evaluate a range of candidate values, specifically $k \in [7, 12]$. This range was selected to ensure a lower bound sufficient for capturing major cell lineages (typically ∼7) while providing an upper bound flexible enough to resolve nuanced, intra-cell-type states through multiple archetypes. For each candidate $k$, we independently generate and align archetypes across modalities, ultimately selecting the value that minimizes the cross-modal discrepancy. For each candidate value $k$, we perform the following:

1. **Assign cells to archetypes.** For each modality $\ell$, each cell $j \in \{1, \ldots, n_\ell\}$ is assigned to its most strongly associated archetype based on its archetype weight vector $s_j^{(\ell)} \in \mathbb{R}^k$ (a row of the reconstruction-weight matrix $S^{(\ell)} \in \mathbb{R}^{n_\ell \times k}$). We denote the archetype assignment and dominance proportion for cell $j$ as:

$$\gamma_j = \arg \max_{i' \in \{1, \ldots, k\}} s_{j,i'}^{(\ell)}, \quad \rho_j = \frac{s_{j,\gamma_j}^{(\ell)}}{\sum_{i'=1}^{k} s_{j,i'}^{(\ell)}} .$$

   For each archetype $i \in \{1, \ldots, k\}$, we define the set of cells assigned to it as:

$$\mathcal{G}_i^{(\ell)} = \{j \in \{1, \ldots, n_\ell\} : \gamma_j = i\} .$$

   These archetype-specific cell sets are used in the subsequent step to characterize each archetype's cell-type composition profile.

2. **Compute cell-type proportion profiles.** For each modality $\ell$, we compute the archetype-by-cell-type proportion matrix $\Phi^{(\ell)} \in \mathbb{R}^{k \times |\mathcal{T}|}$, where each entry $\Phi^{(\ell)}[i, t]$ represents the proportion of cell type $t$ among cells assigned to archetype $i$:

$$\Phi^{(\ell)}[i, t] = \frac{\left|\left\{j \in \mathcal{G}_i^{(\ell)} : c_j = t\right\}\right|}{\left|\mathcal{G}_i^{(\ell)}\right|} ,$$

   where $c_j$ is the known cell type label for cell $j$. This yields a cell-type composition signature for each archetype.

3. **Cross-modal archetype matching.** Given the cell-type proportion matrices $\Phi^{(\mathrm{RNA})} \in \mathbb{R}^{k \times |\mathcal{T}|}$ and $\Phi^{(\mathrm{Prot})} \in \mathbb{R}^{k \times |\mathcal{T}|}$ for RNA and spatial protein modalities, we find the optimal permutation $\pi^*$ that aligns archetypes across modalities by solving the linear assignment problem:

$$\pi^* = \arg \min_{\pi \in S_k} \sum_{i=1}^{k} d_{\mathrm{cosine}} \left(\Phi^{(\mathrm{RNA})}[i, :], \Phi^{(\mathrm{Prot})}[\pi(i), :]\right) ,$$

   where $d_{\mathrm{cosine}}$ is the cosine distance between row vectors, and $S_k$ is the set of all permutations of $k$ elements. This optimization is solved efficiently in $\mathcal{O}(k^3)$ time using the Hungarian algorithm [8].

After evaluating all candidate values, we select the optimal $k^*$ that minimizes the average cross-modal alignment cost (normalized by $k$ to avoid bias toward smaller $k$ values):

$$k^* = \arg \min_{k} \frac{1}{k} \sum_{i=1}^{k} d_{\mathrm{cosine}} \left(\Phi^{(\mathrm{RNA})}[i, :], \Phi^{(\mathrm{Prot})}[\pi^*(i), :]\right) .$$

We then apply the optimal permutation $\pi^*$ to re-index archetypes in each modality, yielding a unified archetype-based coordinate system where corresponding biological archetypes share indices across modalities.

##### Step 3: Unify cell representation across modalities

Each cell is represented as a nonnegative, simplex-constrained mixture of archetypes. For each modality $\ell \in \{\text{RNA}, \text{Spatial Protein}\}$:

$$x^{(\ell)} \approx (A^{(\ell)})^\top s^{(\ell)}, \quad s^{(\ell)} \geq 0, \quad \mathbf{1}^\top s^{(\ell)} = 1,$$

where:

- $x^{(\ell)} \in \mathbb{R}^{d_\ell}$ is the feature vector for a single cell (with $d_\ell \in \{G, P\}$ genes or protein markers),
- $A^{(\ell)} \in \mathbb{R}^{d_\ell \times k}$ is the archetype matrix containing $k$ archetypes as columns, constructed as $A^{(\ell)} = (X^{(\ell)})^\top C^{(\ell)}$ where $X^{(\ell)} \in \mathbb{R}^{n_\ell \times d_\ell}$ is the data matrix (cells as rows) and $C^{(\ell)} \in \mathbb{R}^{n_\ell \times k}$ is the nonnegative coefficient matrix from archetypal analysis,
- $s^{(\ell)} \in \mathbb{R}^k$ is the archetype weight vector for a single cell (a row of $S^{(\ell)} \in \mathbb{R}^{n_\ell \times k}$, where rows are cells and columns are archetype indices).

The archetype weight vectors $s^{(\ell)}$ (or "archetype embeddings") provide modality-agnostic representations. Because archetypes capture mutually apparent biological structure, cells from different modalities with similar weight profiles likely belong to the same underlying cell type or state, regardless of feature correspondence. This allows us to identify and match cells between RNA and protein data based on their archetype similarity rather than direct feature linkage.

##### Step 4: Identify high-confidence anchors

We robustly align modalities by restricting cross-modal matching to high-confidence *anchors* rather than all cells. We identify extreme cell state candidates, termed *anchors*, using cells whose archetype weight vectors are dominated by a single archetype, defined as having one archetype coefficient $\alpha_i > 0.5$. These cells represent the most distinct biological phenotypes in either modality, making them more reliable references for cross-modal matching. The motivation behind anchor selection was supported by testing cross-modal cell-type prediction using nearest neighbors in the shared archetype space—anchors achieved substantially higher F1 scores than non-anchor cells in initial tests.

**Inputs and basic definitions.** Let $S^{(\ell)} \in \mathbb{R}^{n_\ell \times k}$ be the archetype weight matrix for modality $\ell \in \{\text{RNA}, \text{Spatial Protein}\}$. For cell $i$, its weight vector is $s_i^{(\ell)} \in \mathbb{R}^k$ with components $s_{ij}^{(\ell)}$ for $j = 1, \dots, k$. We define dominant archetype labels and the dominance proportion as:

$$y_i = \operatorname*{argmax}_{j \in \{1, \dots, k\}} s_{ij}^{(\ell)}, \qquad p_i = \frac{\max_j s_{ij}^{(\ell)}}{\sum_{j=1}^k s_{ij}^{(\ell)}}.$$

Let $S_c = \{ i \in \{1, \ldots, n_\ell\} : y_i = c \}$ and $n_c = |S_c|$. Let $\mathcal{C}$ be the set of valid archetypes after optional quality filtering ($\mathcal{C} \subseteq \{1, \ldots, k\}$).

**Target percentage and balanced count.** Given a percentage parameter $\beta = 0.5$, the nominal total count for selection is:

$$b_{\text{total}} = \Big\lfloor n_\ell \cdot \beta \Big\rfloor.$$

We distribute this approximately equally across valid archetypes to obtain the per-archetype group count:

$$b = \left\lfloor \frac{b_{\text{total}}}{|\mathcal{C}|} \right\rfloor.$$

Then we apply a per-archetype minimum of 10:

$$m'_c = \max(10, b) \quad \text{for each } c \in \mathcal{C}$$

and use a single balanced cap for all archetypes:

$$m = \min_{c \in \mathcal{C}} m'_c .$$

**Anchor selection.** For each $c \in \mathcal{C}$, select the $m$ cells in $S_c$ with the largest $p_i$. Let the selected set be

$$T_c = \operatorname*{arg\,top}_{i \in S_c}^{m} p_i$$

and define the final anchor cells mask $M \in \{0, 1\}^{n_\ell}$ as

$$M_i = \begin{cases} 1, & \text{if } i \in \bigcup_{c \in \mathcal{C}} T_c, \\ 0, & \text{otherwise.} \end{cases}$$

This method ensures a balanced number of anchors for valid archetypes, avoiding bias toward archetypes with larger numbers of cells.

##### Step 5: Compute cross-modal cell distances in archetype space

For each RNA anchor cell $i \in \mathcal{R}_e$ and protein anchor cell $j \in \mathcal{P}_e$, we compute cosine distance in the shared archetype coordinate space. Let $s_{R,i} \in \mathbb{R}^k$ and $s_{P,j} \in \mathbb{R}^k$ denote the archetype weight vectors (rows of the archetype-weight matrices $S_R$ and $S_P$, respectively). The cosine distance is defined as

$$D^{(\mathcal{A})}_{ij} = 1 - \frac{s_{R,i}^{\top} \, s_{P,j}}{\|s_{R,i}\|_2 \, \|s_{P,j}\|_2}$$

yielding a distance matrix $D^{(\mathcal{A})} \in \mathbb{R}_{\geq 0}^{|\mathcal{R}_e| \times |\mathcal{P}_e|}$ restricted to anchors only. Cells with small archetype-space distances are more likely to be biological counterparts across modalities with similar phenotypes.

##### S2.1 Note on handling multiple batches during archetype analysis

If data from either modality includes multiple experimental batches, archetypes are first learned independently within each batch to avoid filtering out biologically-driven phenotypes that may be batch-specific. Per-batch archetype sets are aligned across batches by computing within-batch archetype-to-cell-type proportion signatures and matching them across batches using cosine distance and optimal linear sum assignment (i.e., Hungarian algorithm). This alignment strategy is analogous to the cross-modal alignment approach in Section S2, but instead yielding batch-agnostic archetype coordinates while preserving batch-specific structures within a given modality.

##### S2.2 Note on cell label resolution specification

ARCADIA's archetype construction and cross-modal alignment rely on mutually represented cell types/states between both datasets. Accordingly, the choice of cell labels inputted into ARCADIA can meaningfully impact alignment and interpretation. Cell annotation should be performed by biological experts for each dataset independently prior to ARCADIA. Consequently, the choice to use rough or fine resolution cell annotations when available (i.e., including only major cell types or additional minor cell states) for ARCADIA is left to the user. This choice largely depends on 1) cell type/state recovery upstream of scRNA-seq (i.e., cell sorting), and 2) confidence in fine-grained annotations from spatial protein marker panels. As these considerations are highly dataset-specific, we encourage users to perform fine-grained cell type annotation and test integration performance with an increasing level of cell label resolution.

#### S3 Detailed Dual Variational Autoencoder Architecture and Training

**Objective and overview.** ARCADIA employs a dual variational autoencoder (VAE) to learn two entangled latent spaces from corresponding scRNA-seq and spatial proteomics data. We train two modality-specific VAEs to reconstruct their inputs (**Figure 1A in the main text**), while aligning latent geometry to preserve cross-modal biological structure from phenotypic variation and spatial cell neighborhood (CN) organization (**Figure 1B in the main text**). Plate model representations of each VAE are depicted in **Figure 1C in the main text**. The dual VAE can enable interpretable cross-modal translation by decoding entangled latent representations in the opposite modality, e.g., predicting counterfactual RNA expression profiles or inferring cell-cell communication based on spatial protein expression. In this study, we assign spatial niche information to scRNA-seq cells to test for spatially-resolved differential gene expression (**Figure 1D in the main text**).

**Network architecture.** Two VAEs are instantiated with modality-specific likelihoods but equivalent latent dimensionality to enable entanglement of the two latent spaces during training and for downstream analysis. The RNA VAE encoder consists of $n_{\text{layers}} = 3$ fully connected layers with $n_{\text{hidden}}^{(\text{RNA})} = 1024$ hidden units per layer, followed by a latent layer of dimension

$n_{\text{latent}} = 60$. The spatial protein VAE encoder consists of $n_{\text{layers}} = 3$ fully connected layers with $n_{\text{hidden}}^{(\text{Prot})} = 512$ hidden units per layer, followed by the same latent dimension $n_{\text{latent}} = 60$. Both encoders use batch normalization and dropout with rate $\rho = 0.1$ for regularization. The decoders mirror the encoder architectures in reverse, mapping from the latent space back to the original feature dimensions.

**Optimization and training configuration.** A single AdamW optimizer manages the parameters of both modality-specific VAEs jointly. The optimizer operates with a learning rate of $\eta = 10^{-3}$, exponential moving average coefficients $(\beta_1, \beta_2) = (0.9, 0.999)$, and decoupled weight decay of $\lambda = 0.001$. Gradients are computed jointly from encoder and decoder parameters of both VAEs, then aggregated in a single backward pass to ensure synchronized updates across modalities. Learning rate scheduling is controlled by `ReduceLROnPlateau`, which monitors the validation loss and reduces the learning rate by a factor of $\alpha = 0.95$ when the loss plateaus for 10 consecutive epochs. Additional hyperparameters for the scheduler include a loss improvement threshold of $\tau_{\text{threshold}} = 0.05$, a minimum learning rate floor of $\eta_{\text{min}} = 10^{-6}$, and a cooldown period of 10 epochs after each reduction before resuming monitoring. To prevent overfitting, training proceeds with independent train/validation splits for each modality (80% training and 20% validation). The validation set is used to monitor model performance on unseen data during training, allowing the `ReduceLROnPlateau` scheduler to monitor validation loss and reduce the learning rate when the validation loss stops improving for 10 consecutive epochs, which helps prevent overfitting by adjusting optimization when the model reaches a performance plateau on unseen data. Gradient clipping at a maximum norm of $\tau = 1.0$ is applied for additional stability. A relatively large batch size of $B = 1024$ is employed to ensure adequate representation of anchors; this is necessary because anchors represent only a small fraction of cells in each modality, and larger batches provide more informative anchors samples for meaningful anchor-guided cross-modal matching loss computation. Models are trained for a maximum of 500 epochs.

**Likelihood functions.** The RNA reconstruction uses a zero-inflated negative binomial (ZINB) likelihood for count data, while the spatial protein reconstruction uses a Gaussian likelihood. To empirically validate these distributional assumptions of the input modalities, we conducted statistical testing on random samples from both the RNA and spatial protein data. For RNA, we sampled genes across a subset of cells, while for spatial protein we sampled features (protein markers concatenated with spatial features) across a subset of cells. For each sampled feature from both modalities, we fit four candidate distributions: Normal, Poisson, Negative Binomial, and Zero Inflated Negative Binomial. We then compared their goodness of fit using the Akaike Information Criterion (AIC), defined as

$$\text{AIC} = 2k - 2\log(\hat{L})$$

where $k$ is the number of parameters in the candidate model and $\hat{L}$ is the maximum likelihood of the fitted model, computed by maximizing the log-likelihood function with respect to the model parameters. For each candidate distribution, parameters were estimated by finding the values that maximize the log-likelihood. The model with the lowest AIC for the majority of sampled features in each modality was selected as the optimal likelihood for reconstruction.

**Model size.** The RNA VAE contains approximately 14.6 million trainable parameters, while the spatial protein VAE contains approximately 1.3 million trainable parameters, for a total of approximately 16 million trainable parameters.

**Batch correction.** Batch labels are incorporated as conditional information into the decoders for each modality, allowing the generative model to absorb residual batch effects in the latent space. Combined with the per-batch archetype alignment, this approach yields consistent latent representations across batches and modalities while preserving biological signal.

**Generative and inference processes.** Assume we have an observed RNA count matrix $x^{\text{RNA}}$ with cells $n = 1..N$ and genes $g = 1..G$ and an observed spatial protein expression matrix $x^{\text{Prot}}$ with cells $m = 1..M$ and features $p = 1..P$. For RNA, we also have observed library size $\ell_n$ provided per cell (e.g., total UMI or size factor), not necessarily modeled as a latent variable. For both $\ell \in \{\text{RNA}, \text{Spatial Protein}\}$ we also consider batch labels $b^{(\ell)} \in \{1, \ldots, K_{(\ell)}\}$ for each cell $n$ or $m$. The generative process is as follows:

For each RNA cell $n$, we draw latent state and batch: $z_n^{\text{RNA}} \sim \mathcal{N}(0, I)$ with batch label $b_n^{\text{RNA}}$. We decode normalized expression $\rho$ and dropout probability $\pi$, and then sample observed RNA counts $x^{\text{RNA}}$ with zero inflation and scale to observed library sizes:

$$\rho_{ng}^{\text{RNA}} = \text{softmax}_g\big(\eta^{\text{RNA}}(z_n^{\text{RNA}}, b_n^{\text{RNA}})\big), \qquad \pi_{ng}^{\text{RNA}} = \sigma\big(\kappa^{\text{RNA}}(z_n^{\text{RNA}}, b_n^{\text{RNA}})\big),$$

$$x_{ng}^{\text{RNA}} \sim \text{ZINB}\Big(\mu_{ng} = \ell_n\, \rho_{ng}^{\text{RNA}},\ \theta_g^{\text{RNA}},\ \pi_{ng}^{\text{RNA}}\Big)$$

where $\eta^{\text{RNA}}(\cdot)$ is a decoder network parameterizing normalized gene expression proportions, $\kappa^{\text{RNA}}(\cdot)$ is a decoder network parameterizing zero-inflation probability, and $\theta_g^{\text{RNA}}$ is gene-level inverse dispersion, all learned during training via gradient descent.

For each spatial protein cell $m$, we draw latent state and batch: $z_m^{\text{Prot}} \sim \mathcal{N}(0, I)$ with batch label $b_m^{\text{Prot}}$. We decode normalized intensity $\iota^{\text{Prot}}$, and then sample observed protein expression $x^{\text{Prot}}$ with Gaussian noice and scale to observed feature expression:

$$\iota_{mp}^{\text{Prot}} = \text{softmax}_p\big(\eta^{\text{Prot}}(z_m^{\text{Prot}}, b_m^{\text{Prot}})\big), \qquad x_{mp}^{\text{Prot}} \mid \iota_{mp}^{\text{Prot}} \sim \mathcal{N}\big(\mu_{mp}^{\text{Prot}} = \alpha^{\text{Prot}}\, \iota_{mp}^{\text{Prot}},\ (\sigma_p^{\text{Prot}})^2\big).$$

where $\eta^{\text{Prot}}(\cdot)$ is a decoder network parameterizing normalized spatial protein feature intensities, $\alpha^{\text{Prot}} > 0$ is a fixed positive hyperparameter scaling decoded normalized intensity to match the observed intensity, and $(\sigma_p^{\text{Prot}})^2$ is the per-feature Gaussian observation noise variance for spatial protein measurements, all learned during training via gradient descent.

For the inference process, we construct variational posteriors for both $\ell \in \{\text{RNA}, \text{S.P.}\}$:

$$q_\phi\big(z_n^{\text{RNA}} \mid x_n^{\text{RNA}}, b_n^{\text{RNA}}\big) = \mathcal{N}\big(\mu_n^{\text{RNA}},\ \text{diag}((\sigma_n^{\text{RNA}})^2)\big),$$

$$q_\phi\big(z_m^{\text{Prot}} \mid x_m^{\text{Prot}}, b_m^{\text{Prot}}\big) = \mathcal{N}\big(\mu_m^{\text{Prot}},\ \text{diag}((\sigma_m^{\text{Prot}})^2)\big).$$

where $\phi$ are neural network parameters of the encoder networks $q_\phi^{\text{RNA}}$ and $q_\phi^{\text{Prot}}$, learned during training via gradient descent, and $\mu$ and $\sigma^2$ are outputs of modality-specific encoder networks that condition on both expression data and batch labels, with $\mu_n^{\text{RNA}} = \ell_n \, \rho_n^{\text{RNA}}$ and $\mu_m^{\text{Prot}} = \alpha^{\text{Prot}} \, \iota_m^{\text{Prot}}$.

**Hyperparameter Selection and Tuning.** Hyperparameters were selected via an automated grid search procedure tracked with MLflow [9]. We explored a parameter space covering network depth (3–5 layers), width (128–1024 units), latent dimensionality (20–60), and loss weighting terms.

We utilized a suite of validation metrics to select the optimal hyperparameters on a held-out set, ensuring the final model balanced integration quality with biological fidelity:

1. **Cross-Modal F1 Score:** Ensures accurate alignment of corresponding cell states across modalities.
2. **iLISI:** Guarantees effective mixing of modalities to prevent technical batch effects.
3. **Moran's I:** Verifies that the latent space preserves the continuous geometry of cell states. High Moran's I ensures a ***smooth phenotypic gradient*** in the latent space, where cells with similar archetype weights (indicating proximity to extreme phenotypic states) cluster together rather than being scattered randomly.
4. **kSep:** Ensures that crucial spatial information is not lost during integration; specifically, it checks that distinct spatial neighborhoods (CNs) remain separable within cell-type clusters.

The final selected architecture employs 3 layers with asymmetric encoder widths ($n_{\text{hidden}}^{\text{RNA}} = 1024$, $n_{\text{hidden}}^{\text{Prot}} = 512$) to account for feature dimensionality difference between modalities, projecting to a shared latent dimension of 60.

To handle the varying scales of different loss components, we implemented an automatic **loss scaling warmup**. During the first 5 epochs, we computed the mean magnitude of each loss term and normalized them such that each component effectively starts with a base weight of 1.0. We then applied specific relative importance weights to these normalized terms. The final configuration used $\lambda_0^{(\text{RNA})} = 1.0$ and $\lambda_0^{(\text{Prot})} = 10.0$ for reconstruction, $\lambda_1 = 0.1$ for structure preservation, $\lambda_2 = 0.1$ for cross-modal alignment, and $\lambda_3 = 1.0$ for anchor matching, as this combination maximized the validation metrics.

#### S3.1 Evidence lower bound (ELBO)

Each VAE optimizes a standard evidence lower bound (ELBO) with modality-specific likelihoods and KL regularization. For a single cell $i$ in modality $\ell$, the ELBO loss is:

$$\mathcal{L}(\theta, \phi; x_i^{(\ell)}, b_i^{(\ell)}) = -\mathbb{E}_{q_\phi(z_i^{(\ell)} | x_i^{(\ell)}, b_i^{(\ell)})} \left[ \log p_\theta(x_i^{(\ell)} \mid z_i^{(\ell)}, b_i^{(\ell)}) \right] + \text{KL}\left( q_\phi(z_i^{(\ell)} \mid x_i^{(\ell)}, b_i^{(\ell)}) \parallel p(z) \right)$$

where:

- $x_i^{(\ell)}$ is the observed feature vector for cell $i$ in modality $\ell$ (RNA gene counts or spatial protein features),
- $b_i^{(\ell)}$ is the batch label for cell $i$ in modality $\ell$ (experimental batch identifier),
- $z_i^{(\ell)}$ is the entangled latent representation of cell $i$ from either VAE,
- $q_\phi(z_i^{(\ell)}|x_i^{(\ell)}, b_i^{(\ell)})$ is the variational encoder mapping observed data to the entangled latent space,
- $p_\theta(x_i^{(\ell)} \mid z_i^{(\ell)}, b_i^{(\ell)})$ is the decoder reconstructing observed data from the latent space.

Training maximizes the ELBO across all cells from both modalities:

$$\mathcal{L}_{\text{ELBO}}^{(\text{RNA})} = -\sum_{n=1}^{N} \Big( \underbrace{\mathbb{E}_{q_\phi(z_n^{\text{RNA}}|x_n^{\text{RNA}})}\big[\log p_\theta(x_n^{\text{RNA}} \mid z_n^{\text{RNA}}, b_n^{\text{RNA}})\big]}_{\text{RNA counts reconstruction (ZINB)}} + \underbrace{\text{KL}\big[q_\phi(z_n^{\text{RNA}} \mid x_n^{\text{RNA}}) \,\|\, p(z)\big]}_{\text{Latent regularization}} \Big),$$

$$\mathcal{L}_{\text{ELBO}}^{(\text{Prot})} = -\sum_{m=1}^{M} \Big( \underbrace{\mathbb{E}_{q_\phi(z_m^{\text{Prot}}|x_m^{\text{Prot}})}\big[\log p_\theta(x_m^{\text{Prot}} \mid z_m^{\text{Prot}}, b_m^{\text{Prot}})\big]}_{\text{Spatial proteomics reconstruction (Gaussian)}} + \underbrace{\text{KL}\big[q_\phi(z_m^{\text{Prot}} \mid x_m^{\text{Prot}}) \,\|\, p(z)\big]}_{\text{Latent regularization}} \Big).$$

$N$ and $M$ are the numbers of RNA and spatial protein cells, respectively (from $X^{(\text{RNA})} \in \mathbb{R}^{N \times G}$ and $X^{(\text{Prot})} \in \mathbb{R}^{M \times P}$). We reconstruct RNA counts using a zero-inflated negative binomial (ZINB) likelihood, and spatial protein expression using a Gaussian likelihood. For both VAEs we employ $p(z) = \mathcal{N}(0, I)$, a standard normal prior on latent representations.

To optimize the ELBO, gradients are estimated via the reparameterization trick for all continuous latent variables, enabling efficient backpropagation through the stochastic latent sampling nodes. This approach allows joint optimization of encoder and decoder parameters across both modalities while maintaining the regularizing effect of the KL divergence terms.

#### S3.2 Anchor-guided cross-modal matching

A matching loss penalizes discrepancies between RNA and spatial protein latent representations within archetypes. Specifically, we use high-confidence anchors to establish cross-modal correspondence before entangling the latent spaces. This loss operates across two distinct spaces for each modality: the archetype embedding space (computed via PCHA) and VAE latent space. The objective is to attract anchors that are nearby in archetype geometry while repelling those that are distant, thereby transferring archetype-derived structure into the VAE latent representations. We use a softplus penalty so that negative margins contribute approximately zero loss while positive margins grow smoothly.

**Sets and data.** Let $\mathcal{R} = \{1, \dots, N_r\}$ index RNA cells and $\mathcal{P} = \{1, \dots, N_{sp}\}$ index spatial protein cells. Let $S_R \in \mathbb{R}^{N_R \times k}$ and $S_P \in \mathbb{R}^{N_P \times k}$ be archetype-weight matrices (where each row is a cell's archetype coordinate vector as defined in Section S2), with rows $s_{R,i}^\top$ and $s_{P,j}^\top$. Let $D \in \mathbb{R}_{\geq 0}^{N_R \times N_P}$ be the matrix of cosine distances in archetype space, with entry $D_{ij}$ for $i \in \mathcal{R}$ and $j \in \mathcal{P}$. Let $\mathcal{R}_e \subseteq \mathcal{R}$ and $\mathcal{P}_e \subseteq \mathcal{P}$ be the anchors sets previously established.

**Cosine geometry in archetype space.** Define row-normalized archetype vectors:

$$\hat{a}_{R,i} \;=\; \frac{a_{R,i}}{\|a_{R,i}\|_2}, \quad i \in \mathcal{R}_e \;, \qquad \hat{a}_{P,j} \;=\; \frac{a_{P,j}}{\|a_{P,j}\|_2}, \quad j \in \mathcal{P}_e \;.$$

Define cosine similarity and distance:

$$S_{ij} \;=\; \hat{a}_{R,i}^{\top}\hat{a}_{P,j}, \qquad D_{ij}^{(\mathcal{A})} \;=\; 1 - S_{ij}, \qquad (i,j) \in \mathcal{R}_e \times \mathcal{P}_e \;.$$

**Anchors latent distances.** Let $D^{(z)} \in \mathbb{R}_{\geq 0}^{N_R \times N_P}$ denote the latent-space distances between RNA and spatial protein cells in the each VAE's latent space. We restrict to anchors:

$$D^{(z,e)} \;=\; \big(D_{ij}^{(z)}\big)_{(i,j)\in\mathcal{R}_e\times\mathcal{P}_e}$$

**Closest and farthest anchors in archetype space.** To align paired anchors across modalities, we identify for each anchor its most similar and most dissimilar counterpart in the opposite modality based on archetype space geometry. This establishes anchor pairs that should be proximal in each latent space (similar archetypes) and contrastive pairs that should be separated (dissimilar archetypes).

For each RNA anchor $i \in \mathcal{R}_e$, define:

$$c^{(R)}(i) \;=\; \arg\min_{j\in\mathcal{P}_e} D_{ij}^{(\mathcal{A})}, \qquad f^{(R)}(i) \;=\; \arg\max_{j\in\mathcal{P}_e} D_{ij}^{(\mathcal{A})},$$

and for each spatial protein anchor $j \in \mathcal{P}_e$, define:

$$c^{(P)}(j) \;=\; \arg\min_{i\in\mathcal{R}_e} D_{ij}^{(\mathcal{A})}, \qquad f^{(P)}(j) \;=\; \arg\max_{i\in\mathcal{R}_e} D_{ij}^{(\mathcal{A})},$$

where $D_{ij}^{(\mathcal{A})} = 1 - \frac{s_{R,i}^{\top}s_{P,j}}{\|s_{R,i}\|_2\|s_{P,j}\|_2}$ is the cosine distance between archetype weight vectors as defined previously. Here, $c^{(R)}(i)$ and $f^{(R)}(i)$ identify the closest and farthest spatial protein anchors to RNA anchor $i$, while $c^{(P)}(j)$ and $f^{(P)}(j)$ identify the closest and farthest RNA anchors to spatial protein anchor $j$.

The corresponding latent distances are extracted from the VAE latent space distance matrix $D^{(z)}$:

$$d_i^{R,\text{close}} \;=\; D^{(z)}_{i,\,c^{(R)}(i)}, \qquad d_i^{R,\text{far}} \;=\; D^{(z)}_{i,\,f^{(R)}(i)}, \qquad i \in \mathcal{R}_e \;,$$

$$d_j^{P,\text{close}} \;=\; D^{(z)}_{c^{(P)}(j),\,j}, \qquad d_j^{P,\text{far}} \;=\; D^{(z)}_{f^{(P)}(j),\,j}, \qquad j \in \mathcal{P}_e$$

These distances measure how well latent spaces preserve archetype-based similarities between anchor pairs.

**Training mini-batch statistics and adaptive margins.** Let

$$\mu \;=\; \frac{1}{|\mathcal{R}_e||\mathcal{P}_e|}\sum_{i\in\mathcal{R}_e}\sum_{j\in\mathcal{P}_e} D_{ij}^{(z)}, \qquad \sigma \;=\; \sqrt{\frac{1}{|\mathcal{R}_e||\mathcal{P}_e|}\sum_{i\in\mathcal{R}_e}\sum_{j\in\mathcal{P}_e}\big(D_{ij}^{(z)} - \mu\big)^2} \;.$$

Define the adaptive margins:

$$m_{\text{close}} \;=\; \alpha_{\text{close}}\,\sigma, \qquad \tau_{\text{far}} \;=\; \alpha_{\text{far}}\,\sigma,$$

where $m_{\text{close}}$ is a small "closeness" margin (with $0 < \alpha_{\text{close}} \ll 1$) and $\tau_{\text{far}}$ is a "farness" margin (with $\alpha_{\text{far}} \geq 1$), ensuring that cells that are close in archetype space remain close in latent space and cells that are far apart remain separated.

**Close pairs.** For closest pairs (similar in archetype space), we penalize only when the latent distance exceeds $m_{\text{close}}$:

$$\text{softplus}\big(d - m_{\text{close}}\big).$$

**Far pairs.** For farthest pairs (dissimilar in archetype space), we penalize only when the latent distance falls below $\tau_{\text{far}}$:

$$\text{softplus}\big(\tau_{\text{far}} - d\big).$$

**Penalties.** With $\text{softplus}_\beta(x) = \beta^{-1}\log(1 + e^{\beta x})$ and $\beta = 0.5$, we calculate

$$\mathcal{L}^{(R)}_{\text{close}} \;=\; \frac{1}{|\mathcal{R}_e|}\sum_{i\in\mathcal{R}_e}\text{softplus}\big(d_i^{R,\text{close}} - m_{\text{close}}\big), \qquad \mathcal{L}^{(P)}_{\text{close}} \;=\; \frac{1}{|\mathcal{P}_e|}\sum_{j\in\mathcal{P}_e}\text{softplus}\big(d_j^{P,\text{close}} - m_{\text{close}}\big),$$

$$\mathcal{L}^{(R)}_{\text{far}} \;=\; \frac{1}{|\mathcal{R}_e|}\sum_{i\in\mathcal{R}_e}\text{softplus}\big(\tau_{\text{far}} - d_i^{R,\text{far}}\big), \qquad \mathcal{L}^{(P)}_{\text{far}} \;=\; \frac{1}{|\mathcal{P}_e|}\sum_{j\in\mathcal{P}_e}\text{softplus}\big(\tau_{\text{far}} - d_j^{P,\text{far}}\big),$$

$$\mathcal{L}_{\text{anchors}} = \mathcal{L}^{(R)}_{\text{close}} + \mathcal{L}^{(R)}_{\text{far}} + \mathcal{L}^{(P)}_{\text{close}} + \mathcal{L}^{(P)}_{\text{far}}\;.$$

#### S3.3 Cell-type structure preservation

This loss preserves within-modality cell-type structure in the latent space. We compute cell-type relationships directly in archetype space and enforce that the VAE latent space preserves this structure.

**Archetype reference.** For each known cell type $t \in \mathcal{T}$ and modality $\ell \in \{\text{RNA}, \text{Spatial Protein}\}$, we first compute centroids in archetype space. Let $S_t \subseteq \{1, \ldots, n_\ell\}$ denote the set of cells belonging to cell type $t$, and let $s_i^{(\ell)} \in \mathbb{R}^k$ denote the archetype weight vector for cell $i$ in modality $\ell$. The centroid for cell type $t$ in archetype space is computed as:

$$\tilde{\mu}_t^{(\ell)} = \frac{1}{|S_t|}\sum_{i\in S_t} s_i^{(\ell)} \in \mathbb{R}^k,$$

where $k$ is the number of archetypes. We construct a reference affinity matrix $\tilde{W}^{(\ell)} \in \mathbb{R}^{|\mathcal{T}|\times|\mathcal{T}|}$ based on inter-type distances in archetype space using a Gaussian kernel:

$$\tilde{W}_{ij}^{(\ell)} = \exp\left(-\frac{\|\tilde{\mu}_i^{(\ell)} - \tilde{\mu}_j^{(\ell)}\|_2^2}{2\sigma_{\text{ref}}^{(\ell),2}}\right) \quad \text{for } i \neq j,$$

where $\tilde{W}_{ij}^{(\ell)}$ represents the affinity (similarity) between cell types $i$ and $j$, and $\sigma_{\text{ref}}^{(\ell)}$ is the standard deviation of pairwise centroid distances in archetype space serving as the Gaussian kernel bandwidth. We employ the Gaussian kernel rather than raw Euclidean distances because archetype and VAE latent spaces may operate at fundamentally different distance scales due to differences in dimensionality, feature ranges, and learned representations. The Gaussian kernel normalizes these distances into a bounded similarity space, enabling meaningful comparison of relative inter-type relationships. Diagonal entries are set to zero to ignore self distance. This reference structure encodes biologically meaningful relationships between cell types and serves as the target geometry for the VAE latent space.

**Structure preservation.** For each known cell type $t \in \mathcal{T}$, we compute centroids in the VAE latent space $z^{(\ell)}$. Let $\mu_t^{(\ell)} \in \mathbb{R}^q$ denote the centroid of all cells of type $t$ in modality $\ell$, where $q$ is the latent dimension. We construct a corresponding affinity matrix $W^{(\ell)} \in \mathbb{R}^{|\mathcal{T}|\times|\mathcal{T}|}$ in latent space using the same Gaussian kernel:

$$W_{ij}^{(\ell)} = \exp\left(-\frac{\|\mu_i^{(\ell)} - \mu_j^{(\ell)}\|_2^2}{2\sigma_\ell^2}\right) \quad \text{for } i \neq j,$$

where $W_{ij}^{(\ell)}$ represents the affinity between cell types $i$ and $j$ in the VAE latent space, and $\sigma_\ell$ is the standard deviation of pairwise centroid distances in latent space. By using the same kernel normalization in both spaces, we can compare structural relationships despite potentially different distance distributions. Structure preservation loss encourages the latent space affinity pattern to match the reference archetype-space pattern:

$$\mathcal{L}_{\text{struct}}^{(\ell)} = \frac{1}{|\mathcal{T}|(|\mathcal{T}|-1)} \sum_{i\neq j} \left(W_{ij}^{(\ell)} - \tilde{W}_{ij}^{(\ell)}\right)^2$$

which minimizes divergence between latent and archetype geometry, ensuring that learned representations preserve the biological structure encoded in the archetype reference.

**Cohesion and separation components.** Within-type cohesion is measured as the average distance from cells to their type centroids, normalized by inter-centroid distances:

$$\mathcal{L}_{\text{cohesion}}^{(\ell)} = \frac{1}{|\mathcal{T}|} \sum_{t\in\mathcal{T}} \frac{\frac{1}{n_t}\sum_{i\in t} \|\mathbf{z}_i^{(\ell)} - \mu_t^{(\ell)}\|_2}{\text{mean}_{\text{inter-centroid}}^{(\ell)}}$$

where $n_t$ is the number of cells of type $t$. Between-type separation encourages type centroids to remain distinct:

$$\mathcal{L}_{\text{sep}}^{(\ell)} = \sum_{i<j} \exp\left(-\|\mu_i^{(\ell)} - \mu_j^{(\ell)}\|_2\right).$$

**Component weighting.** The total cell-type clustering loss combines these components with consistent weights across modalities:

$$\mathcal{L}_{\text{cell-type cluster}}^{(\ell)} = \mathcal{L}_{\text{struct}}^{(\ell)} + \mathcal{L}_{\text{cohesion}}^{(\ell)} + \mathcal{L}_{\text{sep}}^{(\ell)}$$

This design ensures compact, well-separated clusters for each known cell type within each modality's latent space, while maintaining consistency with the reference archetype-derived cell-type structure.

##### S3.4 Cross-modal alignment

To align cell-type representations across modalities, we compute the Maximum Mean Discrepancy (MMD) between latent representations of corresponding cell types in the VAE latent space $z^{(\ell)}$. This loss encourages matching of entire cell-type distributions rather than just centroids.

For each cell type $t \in \mathcal{T}$ present in both modalities, let $Z_t^{(\ell)} \in \mathbb{R}^{N_t^{(\ell)} \times q}$ denote the matrix of latent representations in modality $\ell \in \{\text{RNA}, \text{Spatial Protein}\}$, where $N_t^{(\ell)}$ is the number of type-$t$ cells in modality $\ell$ and $q$ is the latent dimension. Each row of $Z_t^{(\ell)}$ is a single cell's latent embedding. The MMD between the two distributions is computed as:

$$\begin{aligned}\text{MMD}_t^2 = &\frac{1}{N_t^{(\text{RNA})}(N_t^{(\text{RNA})}-1)}\sum_{i\neq j} k(z_i^{(\text{RNA})}, z_j^{(\text{RNA})})\\ &+ \frac{1}{N_t^{(\text{Prot})}(N_t^{(\text{Prot})}-1)}\sum_{i\neq j} k(z_i^{(\text{Prot})}, z_j^{(\text{Prot})})\\ &- \frac{2}{N_t^{(\text{RNA})}N_t^{(\text{Prot})}}\sum_{i,j} k(z_i^{(\text{RNA})}, z_j^{(\text{Prot})})\end{aligned}$$

where the RBF kernel with adaptive bandwidth is:

$$k(x,y) = \exp\left(-\frac{\|x-y\|_2^2}{2\sigma^2}\right)$$

and $\sigma$ is determined adaptively via the median heuristic on combined cross-modal data. Specifically, $\sigma_m = \text{median}\{\|x-y\|_2 : x \in Z_t^{(\text{RNA})}, y \in Z_t^{(\text{Prot})}\}$ automatically scales the kernel to the data's natural distance distribution. Rather than relying on a single bandwidth, an ensemble of MMD values is computed at multiple scales: fine-grained ($\sigma_m^{\text{fine}} = \sigma_m/2$), medium ($\sigma_m^{\text{med}} = \sigma_m$), and coarse-grained ($\sigma_m^{\text{coarse}} = 2\sigma_m$), and averaged equally:

$$\text{MMD}_{t,\text{ensemble}}^2 = \frac{1}{|\Sigma_{\text{scales}}|}\left[\text{MMD}_t^2(\sigma_m^{\text{fine}}) + \text{MMD}_t^2(\sigma_m^{\text{med}}) + \text{MMD}_t^2(\sigma_m^{\text{coarse}})\right]$$

This multi-scale approach provides robustness across varying cell-type geometries: fine-grained scales detect tightly clustered cell types, medium scales capture intermediate structure, and coarse-grained scales capture loosely distributed ones, ensuring the alignment is insensitive to differences in how cell types are dispersed within the latent space.

The cross-modal alignment loss is the average MMD across all common cell types:

$$\mathcal{L}_{\text{cross-modal alignment}} = \frac{1}{|\mathcal{T}_{\text{common}}|}\sum_{t\in\mathcal{T}_{\text{common}}} \text{MMD}_{t,\text{ensemble}}^2$$

where $\mathcal{T}_{\text{common}}$ is the set of cell types present in both modalities, and the subscript "ensemble" denotes the average over the three kernel bandwidths.

**Intuition.** This loss equalizes within-modality distances with cross-modality distances (between cells of different modalities). By matching these distance distributions, intra-modality and cross-modality neighborhoods become indistinguishable. Consequently, the resulting latent space provides no modality structure that could be exploited for clustering cells by measurement platform. Instead, cells organize purely by cell identity, yielding a modality-invariant representation.

##### S3.5 Total loss

**Per-modality total loss.** For each modality $\ell \in \{\text{RNA}, \text{Spatial Protein}\}$, the total loss combines the reconstruction ELBO with geometric constraints:

$$\mathcal{L}^{(\ell)}_{\text{modality}} \;=\; \lambda^{(\ell)}_0 \,\mathcal{L}^{(\ell)}_{\text{ELBO}} \;+\; \lambda^{(\ell)}_1 \,\mathcal{L}^{(\ell)}_{\text{cell-type cluster}} \tag{S1}$$

where $\lambda^{(\ell)}_0, \lambda^{(\ell)}_1$ are hyperparameters controlling the relative importance of each term for modality $\ell$.

**Combined objective.** The overall training objective combines contributions from both modalities and adds the cross-modal losses:

$$\mathcal{L}_{\text{total}} \;=\; \mathcal{L}^{(\text{RNA})}_{\text{modality}} + \mathcal{L}^{(\text{Prot})}_{\text{modality}} + \lambda_2 \,\mathcal{L}_{\text{anchors}} + \lambda_3 \,\mathcal{L}_{\text{cross-modal alignment}} \;, \tag{S2}$$

where $\lambda_2 > 0$ and $\lambda_3 > 0$ control the strength of cross-modal losses.

**Loss scaling via gradient magnitude estimation.** To ensure balanced contributions across loss components with different intrinsic scales, we employ an automatic, gradient-based warmup phase before standard model training. During this phase, we freeze all model parameters (no weight updates) and run $n_{\text{warmup}} = 5$ epochs with all loss weights initialized to $\lambda_0 = \lambda_1 = \lambda_2 = \lambda_3 = 1.0$. For each loss component $\mathcal{L}_i$ with $i \in \{0, 1, 2, 3\}$, we record the loss values (and corresponding gradients) at each warmup epoch without updating the network. We then compute the median loss for each component:

$$\tilde{\mathcal{L}}_i = \text{median}\{\mathcal{L}^{(1)}_i, \mathcal{L}^{(2)}_i, \ldots, \mathcal{L}^{(n_{\text{warmup}})}_i\},$$

and define the loss weights as:

$$\lambda_i = \frac{h}{\tilde{\mathcal{L}}_i},$$

where $h$ is a global reference scale. This procedure automatically rescales each term so that all losses enter the combined objective on comparable magnitude. After this initialization phase, standard training resumes with the $\lambda_i$ values fixed and full model parameter updates enabled. In practice, this auto-calibration yields similar but not identical $\lambda_i$ across datasets, without requiring any manual tuning of loss weights.

#### S4 Baseline Method Implementation Details

Complete reproduction code is available at https://github.com/azizilab/arcadia_reproducibility.

**We selected MaxFuse and scMODAL as the primary benchmarks because they represent the current state-of-the-art for *weak-linkage* integration scenarios, where modalities share limited or noisy feature sets** (unlike totalVI which requires paired barcodes, or BindSC/SCOT which struggle with disparate feature spaces). See **Section S6** for additional discussion and comparisons with state-of-the-art integration methods.

To generate the benchmarking metrics presented in the Results section of the main text, we executed the baseline methods using their official Python implementations with the following protocols:

- **MaxFuse [10]:** We utilized the `maxfuse` package. The model was run using the default iterative matching procedure. We used the standard pivot-gene selection strategy (identifying shared highly variable genes) and initialized the alignment using Canonical Correlation Analysis (CCA) as recommended in the documentation.
- **scMODAL [11]:** We utilized the official GitHub repository implementation. The model was trained using the default adversarial hyperparameters. For fair comparison, we provided scMODAL with the same preprocessed gene and protein matrices used for ARCADIA, allowing it to leverage its required feature-matching priors.

For both baselines, we extracted the final latent embeddings to compute the integration metrics (including iLISI, Silhouette, and kSep) and cell-type prediction scores reported in Table 1 in the main text.

#### S5 Detailed Benchmarking Metrics for Semi-Synthetic and Tonsil Datasets

**Standard Evaluation Metrics** To benchmark the performance of ARCADIA, we employ several standard integration evaluation metrics previously used by state-of-the-art methods [10, 11], which quantify: cell type classification accuracy (F1 score [12]), clustering (Adjusted Rand Index (ARI) [13] and silhouette score [14]), and integration and cell mixing (integration Local Inverse Simpson's Index (iLISI) [15] and k-nearest-neighbor batch-effect test (kBET) [16]). To ensure robustness, each metric was computed across 10 independent runs of the ARCADIA integration framework.

**Moran's I.** Moran's I is a spatial autocorrelation statistic that measures whether similar score values cluster together spatially in the embedding space. In this scenario, we implement Moran's I to assess the degree to which cells with similar archetype weights are located near each other in the latent representation.

The metric is computed using the formula:

$$I = \frac{n}{S_0} \cdot \frac{\sum_{i,j} W_{ij}(x_i - \bar{x})(x_j - \bar{x})}{\sum_i (x_i - \bar{x})^2}$$

Table S1: Integration Performance Benchmarking Across Datasets

| | Semi-Synthetic CITE-seq Dataset | | | Tonsil Dataset | | |
|---|---|---|---|---|---|---|
| Metric | ARCADIA | MaxFuse | scMODAL | ARCADIA | MaxFuse | scMODAL |
| kSep | **0.666 ± 0.046** | 0.403 | 0.352 | **0.510 ± 0.016** | 0.212 | 0.449 |
| Moran's I | **0.570 ± 0.041** | 0.395 | 0.493 | **0.606 ± 0.021** | 0.469 | 0.436 |
| Pair Distance† | **0.348 ± 0.069** | 0.850 | 0.640 | – | – | – |
| CN F1 | **0.979 ± 0.003** | 0.556 | 0.577 | – | – | – |
| Cell-type F1 | 0.941 ± 0.011 | 0.921 | **0.963** | **0.897 ± 0.003** | 0.888 | 0.848 |
| ARI Score | 0.972 ± 0.008 | 0.955 | **0.991** | **0.791 ± 0.011** | 0.680 | 0.635 |
| iLISI | **1.771 ± 0.050** | 1.704 | 1.600 | 1.543 ± 0.038 | 1.208 | **1.690** |
| Silhouette Score | **0.607 ± 0.028** | 0.086 | 0.301 | **0.315 ± 0.017** | 0.061 | 0.159 |
| Silhouette F1 | **0.616 ± 0.004** | 0.519 | 0.564 | **0.568 ± 0.004** | 0.520 | 0.538 |
| kBET† | 0.524 ± 0.064 | 0.401 | **0.347** | 0.477 ± 0.018 | **0.246** | 0.584 |

Bold values indicate best performance on each metric. † indicates lower is better. ARCADIA results are presented as mean ± standard deviation across 10 independent runs. "–" indicates that the metric was not applicable to the dataset.

where $n$ is the number of cells, $W_{ij}$ is the spatial weight matrix based on k-nearest neighbors in the embedding space, $x_i$ are the archetype weight scores, $\bar{x}$ is the mean score, and $S_0 = \sum_{i,j} W_{ij}$ is the sum of all spatial weights.

The statistic ranges approximately from $-1$ to $+1$. $I \approx -1$ indicates cells with similar archetype weight scores are distant in the embedding space, and $I \approx 0$ indicates no autocorrelation (archetype weights are randomly distributed with no embedding pattern). $I \approx +1$ reflects that cells with similarly high archetype weights are closer together in the embedding space. Thus, more positive Moran's I values indicate better retention of archetype geometry in the latent space.

**kSep (Per-Cell-Type kBET).** kSep is a cell-type-stratified variant of the k-Batch Effect Test (kBET) [16] and evaluates whether spatial niches segregate within cell type clusters in the integrated latent space. For each cell type cluster, kSep uses $\chi^2$ statistics to test whether the cellular neighborhood (CN) composition around each cell is well-mixed. We use this metric to evaluate to what degree latent space representations reflect biological differences due to spatial context. Higher kSep values suggest latent factors, which inherently encode phenotypic variation within cell types from gene expression, also reflect spatially-dependent cell states captured in spatial protein data.

**Pair Distance.** For multi-modal datasets with paired barcodes across modalities (e.g., CITE-seq), the Pair Distance metric quantifies how well the integrated latent representation preserves the cell-to-cell correspondence between paired cells. Lower values indicate better integration.

Let $m \in \{\text{RNA}, \text{Protein}\}$ denote the two modalities, with paired cells indexed by $i \in \{1, \ldots, n_{\text{pairs}}\}$. Let $\mathbf{z}_i^m \in \mathbb{R}^q$ denote the latent embedding for cell $i$ in modality $m$, where $q$ is the latent dimensionality. For each pair $(i, j)$ representing the same cell measured in different modalities, the *true pair distance* is:

$$d_{\text{pair}}(i, j) = \|\mathbf{z}_i^{\text{RNA}} - \mathbf{z}_j^{\text{Prot}}\|_2$$

The cross-modal neighborhood distances represent all Euclidean distances between randomly paired cells from different modalities:

$$\mathcal{D}_{\text{cross}} = \{d_{ik} = \|\mathbf{z}_i^{\text{RNA}} - \mathbf{z}_k^{\text{Prot}}\|_2 : i \in \text{RNA}, k \in \text{Prot}\}$$

The Pair Distance metric then normalizes the mean true pair distance by the mean cross-modal distance:

$$\text{Pair Distance} = \frac{\text{mean}(d_{\text{pair}})}{\text{mean}(\mathcal{D}_{\text{cross}})}$$

### S6 Comparison with State-of-the-Art Integration Methods

To contextualize ARCADIA's performance, we compared it against leading unsupervised integration methods that do not require paired cells. However, each of these methods relies on distinct algorithmic assumptions, ranging from linear correlation to topological matching, that limit their applicability to spatially-resolved proteomics integration.

**BindSC** utilizes a bi-order Canonical Correlation Analysis (Bi-CCA) to bridge modalities. It works by iteratively optimizing a transition matrix that defines cell-to-cell correspondences while simultaneously identifying correlated features [17]. While effective, BindSC's computational complexity scales as $O(N \cdot p^2 + p^3)$, where $N$ is the number of cells and $p$ is the number of features. The cubic dependency on features ($p^3$) during eigendecomposition makes it computationally expensive for high-dimensional scRNA-seq integration without aggressive feature selection, while the $N$ dependency ensures costs rise with dataset size. In contrast, ARCADIA employs deep neural networks (VAEs) to capture complex, non-linear dependencies between transcriptomic and proteomic profiles with linear scalability. Furthermore, while BindSC infers correspondence through iterative optimization of feature correlations, ARCADIA anchors integration on "phenotypic archetypes," providing a robust signal for alignment even when direct feature correlations are weak.

**SCOT (Single-Cell Optimal Transport)** applies Gromov-Wasserstein optimal transport to align datasets based on their internal geometry [18]. It operates by matching intra-dataset distance matrices to align global topologies. A primary challenge with standard optimal transport formulations is the assumption of mass conservation, which can force global probability matching even when cell type compositions are unbalanced. While recent extensions (e.g., semi-coupled OT) attempt to mitigate this, they introduce complexity in balancing transport cost against mass relaxation terms, often requiring careful hyperparameter tuning. ARCADIA naturally handles unbalanced compositions by anchoring on shared phenotypic extremes (archetypes) rather than forcing a global distributional match. Additionally, while approximation strategies using **entropy-regularized Sinkhorn iterations** operate with a complexity of roughly $O(N \log N)$, the global nature of the optimization often remains more memory-intensive than ARCADIA's pure mini-batch processing.

**UnionCom** performs unsupervised topological alignment by embedding the intrinsic low-dimensional structures of each dataset into distance matrices and aligning them via generalized manifold alignment [19]. Like SCOT, it relies heavily on the assumption that the global topology is preserved across modalities. This poses a challenge in spatial proteomics, where limited panel sizes (e.g., $< 50$ proteins markers) often result in a sparse manifold that lacks the topological fidelity of the whole-transcriptome space. ARCADIA addresses this by injecting "phenotypic distinctness" as a guiding signal; by identifying archetypes, it creates high-confidence anchors that persist even in sparse protein manifolds, ensuring accurate alignment where purely topological methods might struggle with manifold distortion.

**MaxFuse** represents a category of iterative matching algorithms that rely on prior biological knowledge of "feature links" to bridge modalities [10]. It constructs a shared embedding by first identifying pivot features common to both datasets (e.g., homologous genes and proteins) and then iteratively refining cell-to-cell matches. MaxFuse utilizes iterative refinement steps involving nearest-neighbor approximations, scaling effectively as $O(N \log N)$. ARCADIA on the other hand does not make any "feature links" assumption and use the data to direct the relationship between he features of the two modalities.

**scMODAL** represents a class of deep learning methods that utilize Generative Adversarial Networks (GANs) for integration [11]. Like ARCADIA, it offers linear scalability ($O(N)$). However, scMODAL is explicitly designed to leverage "feature links", or prior knowledge of correlations between specific genes and proteins, to seed its alignment (similar to MaxFuse). While powerful when such links are known, this dependency can limit utility in discovery scenarios using novel protein panels where correlations are unestablished. ARCADIA offers a complementary approach by learning a shared latent space via archetype alignment.

Other deep generative models like **totalVI** [20] are designed for multi-modal integration where modalities are linked via **paired cell barcodes**. By leveraging these shared identifiers, totalVI trains on cells where both transcriptomic and proteomic profiles are simultaneously observed, allowing it to learn a joint probabilistic representation $p(x_{\text{RNA}}, x_{\text{Prot}} \mid z)$. However, this reliance on *strong linkage* renders totalVI unsuitable for the integration of dissociated scRNA-seq and spatial proteomics. In contrast, ARCADIA integrates data collected from distinct sets of cells without shared barcodes and from modalities with varying dimensionality. Consequently, standard multi-modal VAEs cannot learn the joint distribution required to bridge these modalities, necessitating a method like ARCADIA that aligns data based on shared biological structure rather than cell-level correspondence.

#### S6.1 Summary of ARCADIA's Contribution

ARCADIA distinguishes itself as a method that combines the non-linear modeling capacity of deep learning with the robustness of archetype-based anchoring. It offers an alternative to the linearity of BindSC, the topological constraints of SCOT and UnionCom, and the feature-prior reliance of MaxFuse and scMODAL. It is important to note that while ARCADIA avoids explicit feature priors, it relies on the assumption that **phenotypic extremes (archetypes) are conserved across modalities**. Uniquely, ARCADIA's generative VAE architecture enables bidirectional translation, extending its utility beyond integration space coordinate alignment.

Table S2: Comparison of ARCADIA with Existing Integration Methods

| Method | Core Algorithm | Alignment Strategy | Handling of Weak/No Links | Robustness to Unbalanced Data | Algorithmic Complexity |
|---|---|---|---|---|---|
| **ARCADIA** | Dual VAE | Aligns **Archetypes** (Phenotypic Extremes) | **Robust:** Uses archetypes as anchors; ignores misleading topology | **High:** Anchors via archetypes; no mass conservation constraint | **Linear** $O(N)$ (Mini-batch SGD) |
| **MaxFuse** [10] | Iterative Matching | Aligns Weak Features (Pivots) | **Good:** Requires known shared features (Gene $\approx$ Protein) | **High** | **Iterative** $O(N \log N)$ |
| **scMODAL** [11] | Deep Learning | Adversarial Alignment | **Good:** Requires known shared features (Gene $\approx$ Protein) | **Medium** | **Linear** $O(N)$ (Mini-batch) |
| **BindSC** [17] | Bi-order CCA | Simultaneous Feature/Cell Alignment | **Risky:** Assumes **Linear** correlation exists | **Medium** | **Iterative** $O(N \cdot p^2 + p^3)$ |
| **SCOT** [18] | Optimal Transport | Global Topology (Shape Matching) | **Risky:** Ignores features; protein/RNA shapes must match | **Low:** Standard OT assumes balanced mass | **Approx** $O(N \log N)$ (Sinkhorn) |
| **UnionCom** [19] | Matrix Optimization | Global Topology (Manifold Alignment) | **Risky:** Ignores features; sensitive to manifold distortion | **Low** | **Approx** $O(N \cdot d^2)$ (Low-rank) |

#### S6.2 Computational Resource Usage (Theoretical Analysis)

*Note: As empirical runtime and memory measurements for baselines were not collected in this study, we compare the theoretical scalability of the underlying algorithms.*

The fundamental difference in computational scalability arises from the optimization strategies employed:

1. **Deep Learning Methods (ARCADIA & scMODAL):**
   - **Scalability:** $O(N)$. These methods utilize stochastic gradient descent with mini-batches. This decouples memory usage from the total number of cells ($N$).

- **Implication:** Memory consumption remains constant (dependent on batch size and model depth) even as the dataset grows to millions of cells.

2. **Global Matrix Methods (SCOT, UnionCom):**

   - **Scalability:** Typically $O(N^2)$ to $O(N^3)$ for naive implementations involving full pairwise distance matrices or global matrix operations.
   - **Implication:** While approximation techniques (e.g., Nyström method, **entropy-regularized Sinkhorn iterations**) can reduce this complexity, the global nature of the optimization often requires significantly higher memory and compute time compared to mini-batch approaches as $N$ increases, potentially requiring downsampling for very large spatial maps.

3. **Feature-Dense Methods (BindSC):**

   - **Scalability:** $O(N \cdot p^2 + p^3)$. The complexity involves both cell-dependent iterations and feature-dependent covariance matrix operations in CCA.
   - **Implication:** While efficient for datasets with few features (e.g., protein-only), runtime can degrade significantly when integrating high-dimensional scRNA-seq data without aggressive feature selection.

### S7 ARCADIA Runtime and Computational Efficiency

To assess the computational requirements and scalability of ARCADIA, we benchmarked the complete integration pipeline on two datasets of varying size and dimensionality: the semi-synthetic CITE-seq dataset (15,337 RNA cells × 1,796 genes; 14,021 protein cells × 220 markers) and the larger human tonsil dataset (12,917 RNA cells × 2,000 genes; 169,468 protein cells × 92 markers). Benchmarks were conducted on a Linux-based server (kernel 5.10.0-37) equipped with an Intel Xeon CPU (8 physical/16 logical cores at 2.20 GHz), 58.87 GB RAM, and a single NVIDIA Tesla T4 GPU (15 GB VRAM).

The pipeline comprises two distinct computational phases with differing complexity characteristics. The **preprocessing phase** (CPU-intensive) encompasses spatial graph construction, normalization, and archetype generation via PCHA. This phase scales super-linearly, dominated by the $k$-nearest neighbor search required for spatial graphs and the iterative convex hull optimization. Consequently, the CITE-seq dataset (∼30k total cells) required 13.5 minutes (14.9 GB peak RAM), while the significantly larger tonsil dataset (∼180k total cells) required 56.9 minutes (38.6 GB peak RAM).

The **dual-VAE training phase** (GPU-accelerated) scales linearly $O(N)$ with respect to cell count due to mini-batch stochastic gradient descent. Training times were comparable across datasets despite the six-fold difference in total cell numbers: 1.3 hours for CITE-seq and 1.8 hours for the tonsil dataset (300 epochs). This efficiency arises because the computational cost per batch is largely determined by the neural network architecture dimensions rather than the total dataset size. Both models utilized similar capacities (∼15.3 million parameters for CITE-seq vs. ∼16.0 million for tonsil) resulting in stable iteration speeds.

Memory profiling indicates that ARCADIA is well-optimized for consumer-grade hardware regarding GPU usage, though system RAM requirements scale with dataset size. Using a batch size of 1,024, peak GPU VRAM utilization was consistently low (3.0 GB for CITE-seq, 3.2 GB for tonsil), suggesting compatibility with GPUs possessing as little as 8 GB VRAM. However, peak System RAM usage during training differed significantly: 7.7 GB for CITE-seq versus 41.7 GB for the tonsil dataset, reflecting the memory cost of loading the larger protein matrix into the dataloader. Furthermore, the pipeline effectively leverages multi-core architectures, evidenced by average CPU utilizations of 196% and 189% (relative to single-core performance) during training.

Crucially, this computational efficiency does not come at the cost of performance. Within these training windows, the model converged to high degrees of cross-modal alignment, achieving training cell-type matching accuracies of 98.3% (CITE-seq) and 88.5% (tonsil), with corresponding iLISI scores of 1.89 and 1.80, indicating effective mixing of modalities in the latent space. With total pipeline runtimes of under 3 hours for large-scale atlases, ARCADIA demonstrates practical scalability for spatial multi-omics integration.

### S8 Detailed Analysis of Spatially-Resolved Gene Expression Patterns in the Tonsil Dataset

As summarized in the main text, we performed differentially expressed gene (DEG) analysis within cell types but across inferred cell neighborhoods (CNs) in the human tonsil dataset. DEG testing was restricted to CNs with adequate cell type representation (**Figure S3A**). Complete lists of the top 50 upregulated DEGs are shown in **Figures S4, S5, and S6**.

#### S8.1 B cells

As a sanity check, we first examined canonical B cell behaviors [21, 22, 23] across germinal center (GC) subniches, represented by CN_3, CN_7, and CN_9 in the dataset (**Figure 5A in the main text**). Top upregulated B cell DEGs in CN_3 reflect somatic hypermutation (*BCL6, AICDA, MYBL1, BACH2, ZBTB20, BRWD1, BCL7A*), B cell activation and B cell receptor (BCR) signaling (*CD84, MME, LRMP, KLHL6, CXCR4*), and proliferation (*MKI67, BIRC5*) in B cells. In comparison, B cells in CN_7 and CN_9 upregulated genes associated with plasma cell differentiation (*CD83, FCRL5*), survival and metabolic remodeling (*GAPDH, LDHA, ODC1*), and immune presentation (*HLA-A, HLA-DQA1, HLA-DQB1*), in addition to activation and signaling (*LCP1, PLEK*) and cell cycle progression (*MYBL2, BIRC5*).

Taken together, ARCADIA uncovers that within human tonsils, B cells residing in CN_3, which often appears at the core of GCs, appear to proliferate highly and undergo somatic hypermutation to promote BCR diversity, whereas B cells in distal GC regions CN_7 and CN_9 may more readily differentiate into immune-surveilling, antibody-secreting plasma cells. These biological insights build upon previous studies, as while they predict RNA expression differences between B cells inside and outside GC boundaries by comparing matched subtypes [10] or describe broad pathways such as G2M checkpoint, TNFA signaling, IL-2 STAT5 signaling, and interferon-gamma response within zones of GCs [24], they do not report phenotypic heterogeneity of functional markers directly reflecting proliferation, selection, and antigen presentation in the same B cell subtype across GC zones.

##### S8.2 T cells

Next, we studied the spatially-dependent phenotypic behavior of CD8 T (**Figure 5B in the main text**) and CD4 T cells (**Figure 5C in the main text**) in neighborhoods surrounding germinal centers, which had not been previously characterized for this human tonsil dataset. In particular, we focus on CD8 or CD4 T cells from CN_0, CN_1, CN_4, CN_5, and CN_8, which represent pockets of immune cells surrounding the germinal centers. While the role of CD4 T cells, specifically follicular helper CD4 T subtypes (Tfh), in GC formation and maintenance is well documented [25], the role of follicular CD8 T cells in relation to GCs has only very recently been examined in depth [26].

CN_0 CD8 T cells upregulate cytotoxic effector (*TNFRSF4, TNFRSF18, TNFRSF25, CTSW*), activation (*FOS, CD7, LTB, IER2, FOSB*), and, notably, stem-like (*IL7R, SELL, TCF7*) markers. In CN_8, and to a lesser extent in CN_1, CN_4, and CN_5, CD8 T cells similarly upregulate cytotoxic effector (*CCL4, GZMA, CCL5, SH2D1A, GZMK, GZMM, CST7, NKG7, EOMES*) and activation (*ITM2A, CD6, SLAMF7, CD84*) markers, but also T cell receptor (*GRAP2, TRAT1, TRAC, TRBC2, FYN*), antigen-presentation (*HLA-A*), and exhaustion (*TIGIT, KLRG1, PDCD4*) markers.

CN_4 is markedly characterized by terminally differentiated (*ZEB2*), highly exhausted CD4 T cell states (*LAG3, PDCD1, CTLA4, CD200, TIGIT, TOX2*). Conversely, CD4 T cells in CN_1 exhibit exhausted but metabolically active and antigen-presenting Tfh phenotypes (*BCL6, MAF, GAPDH, HLA-A*), while CN_5 and CN_8 both feature stem-like CD4 T cells (*CCR7, IL7R*) possibly undergoing further differentiation (*GIMAP1, GIMAP7, GIMAP4*). While the exact role of follicular CD8 T cells, and the importance of the signals they produce, requires additional elucidation, understanding spatially-dependent patterns of CD4 Tfh activation and exhaustion programs is crucial to promote successful CD40-CD40L signaling, B cell activation, and protective immunity, e.g., rational vaccine design, or to better profile and design treatments for poorly managed autoimmune diseases involving derailed B cell response, e.g., rheumatoid arthritis and systemic lupus erythematosus [25].

In sum, ARCADIA improves upon previous methods that alone cannot predict phenotypic heterogeneity as a function of spatial context at a single-cell level, and in doing so it reveals coordinated transcriptional programs of B cell maturation and T cell activation/exhaustion that are dependent on cellular location relative to germinal centers in human tonsils, which have meaningful implications for enhancing vaccine development or treating autoimmunity. These results confirm ARCADIA's utility in synthesizing and linking biologically meaningful spatial and transcriptomic patterns of cellular behavior at the single-cell level.

### S9 Supplemental Figures

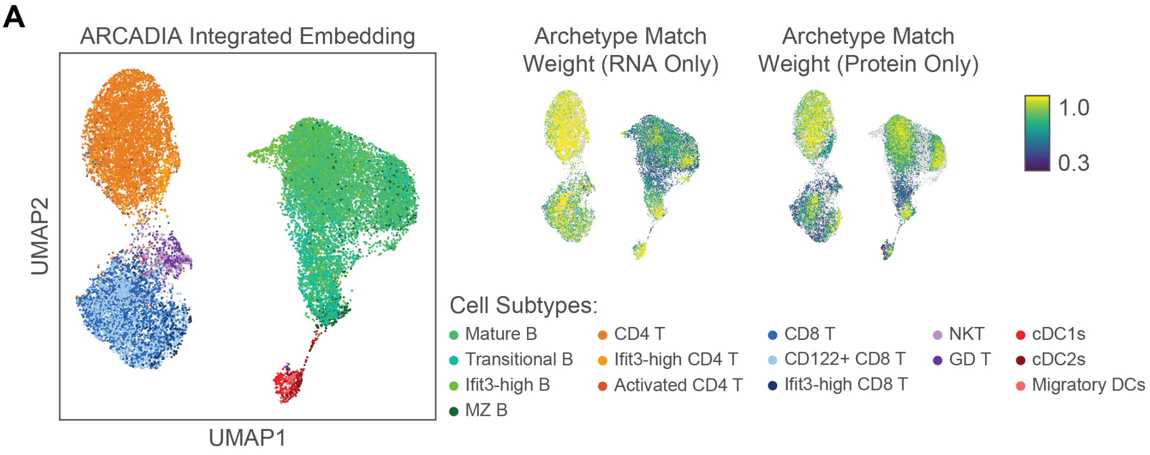


Figure S1: **Additional materials on ARCADIA archetype matching and integration framework.** **(A)** ARCADIA integrated embedding of protein and RNA cells colored by cell subtypes (left), and archetype match weight for cells in RNA and protein data (right).

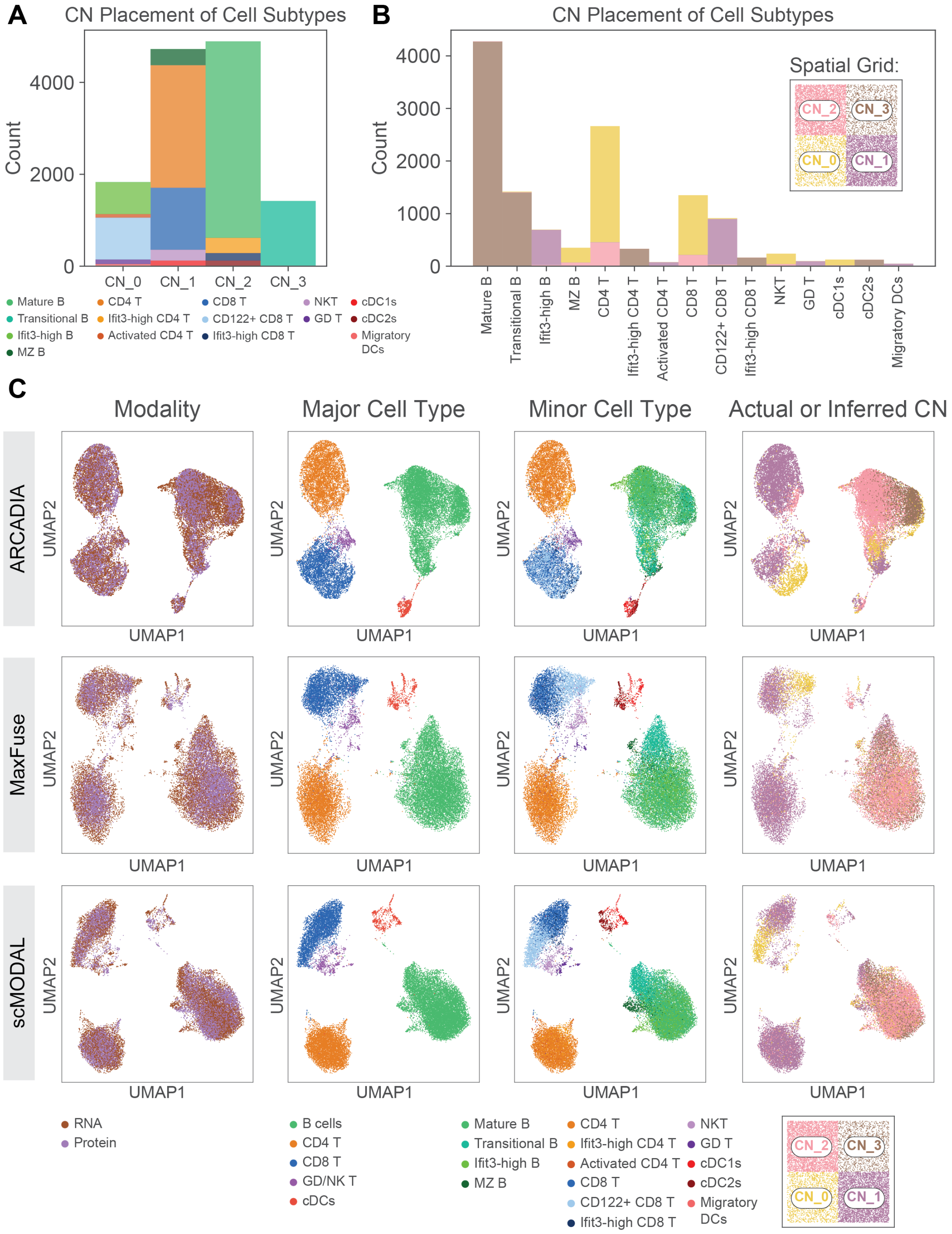


Figure S2: **Additional materials on semi-synthetic data setup, ARCADIA integration, and integration comparison between ARCADIA and MaxFuse and scMODAL.** **(A)** Cell subtype composition of each cell neighborhood in the semi-synthetic spatial grid of protein cells. **(B)** Cell neighborhood membership for each cell subtype in the semi-synthetic spatial grid. **(C)** Integration of protein and RNA cells by ARCADIA (top), MaxFuse (center), and scMODAL (bottom) colored by modality, cell type, subtype, and CN (ground truth for protein, inferred by respective methods for RNA). CN: Cell Neighborhood.

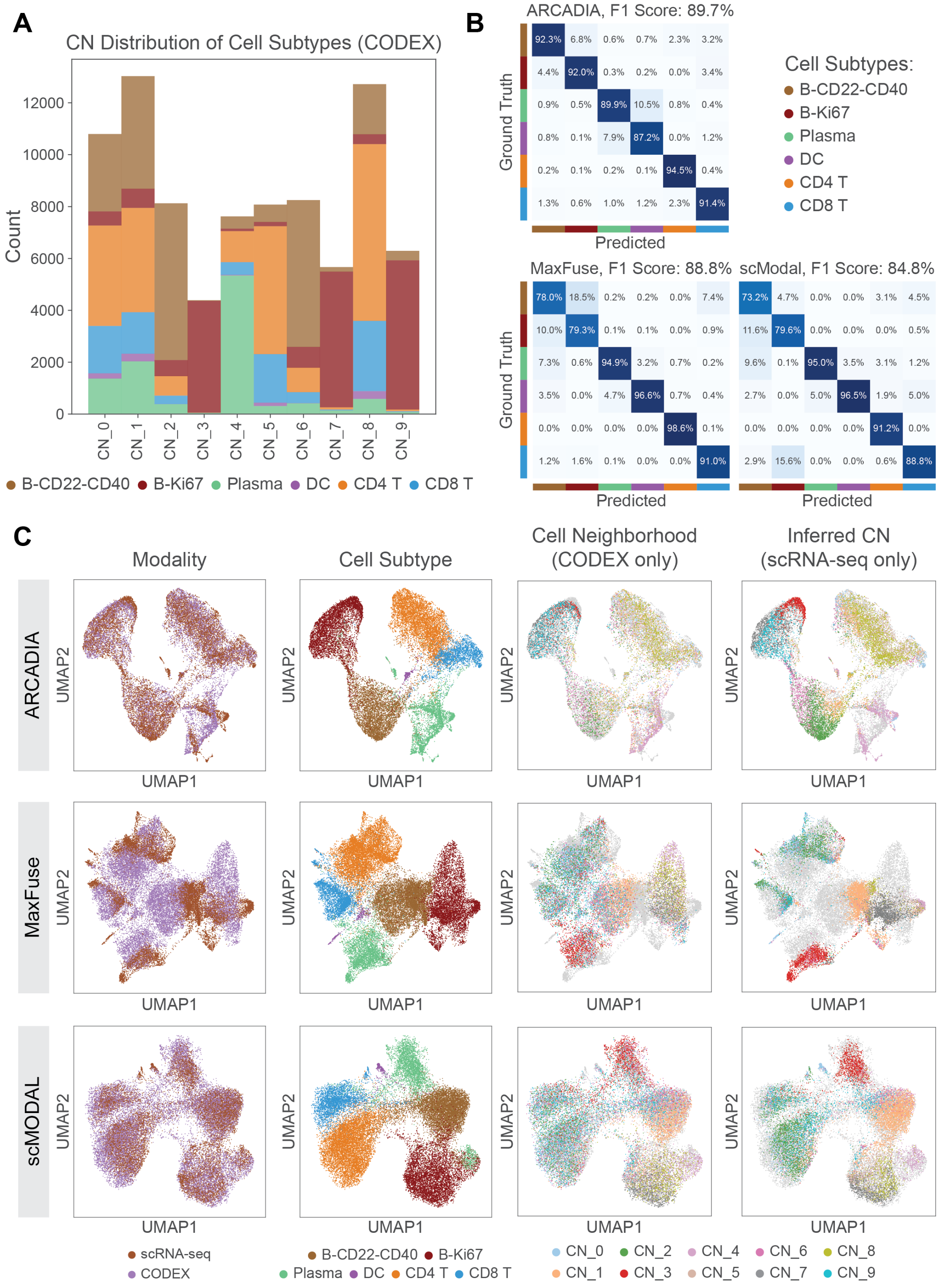


Figure S3: **Additional materials on human tonsil data, ARCADIA integration, and benchmarking. (A)** Cell type composition of each CN in the CODEX data. **(B)** Cell type matching accuracy for ARCADIA, MaxFuse, and scMODAL. **(C)** Integration of protein and RNA data by ARCADIA (top), MaxFuse (center) and scMODAL (bottom) colored by modality, cell type, and ground truth or inferred CN.

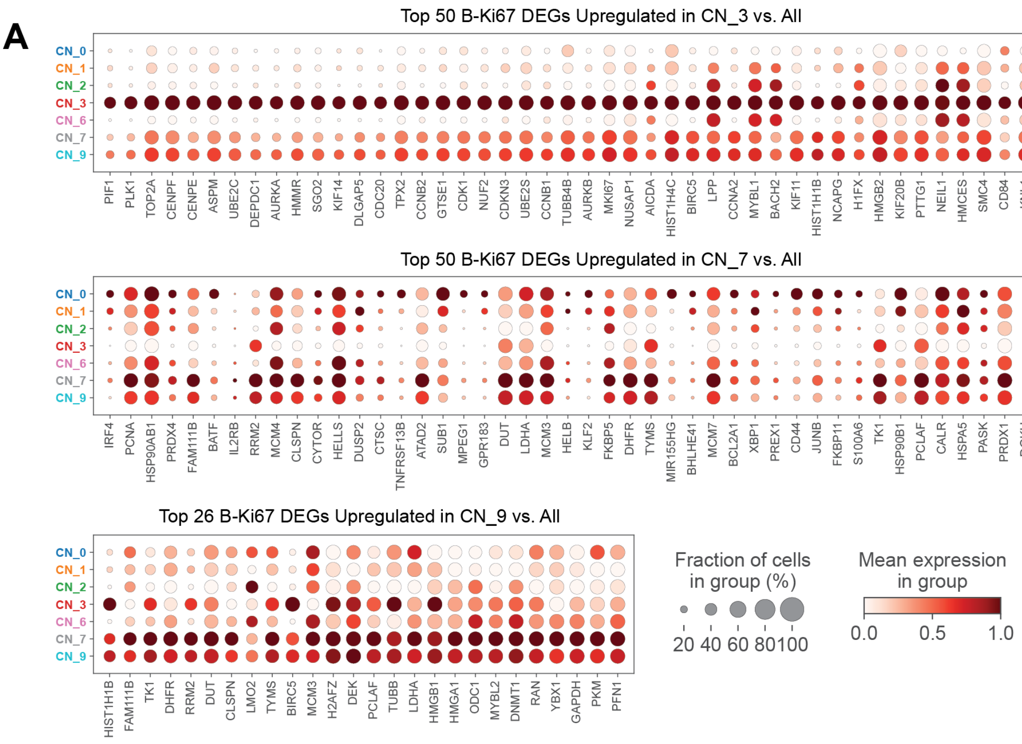


Figure S4: **Additional materials on spatially-resolved differential expression within B-Ki67 cell populations. (A)** Top upregulated differentially expressed genes (DEGs) and their expression between B-Ki67 cells from scRNA-seq predicted to originate from a given cell neighborhood compared to all other B-Ki67 cells. Only cell neighborhoods with ample cell type representation are shown (see cell type composition plot in Figure S3A for more details). Expression for each gene is relative across the comparison groups shown.

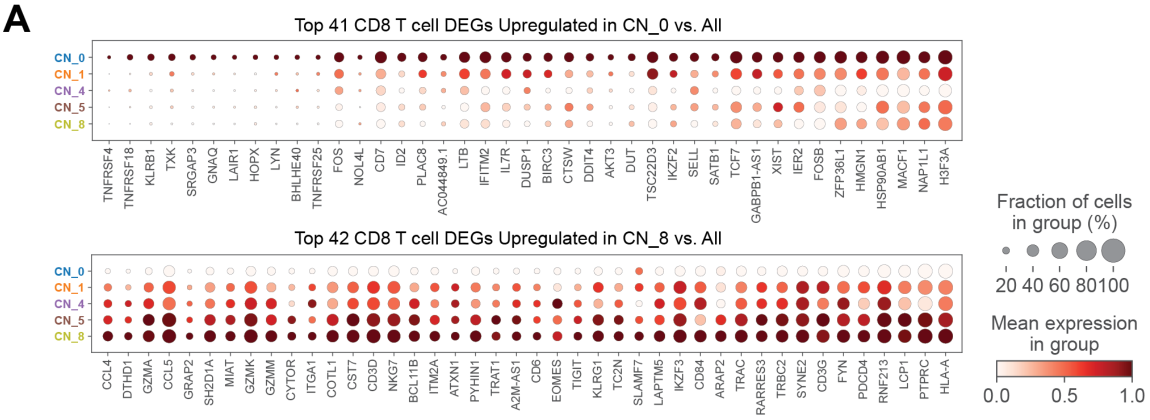


Figure S5: **Additional materials on spatially-resolved differential expression within CD8 T cell populations. (A)** Top upregulated differentially expressed genes (DEGs) and their expression between CD8 T cells from scRNA-seq predicted to originate from a given cell neighborhood compared to all other CD8 T cells. Only cell neighborhoods with ample cell type representation are shown (see cell type composition plot in Figure S3A for more details). Expression for each gene is relative across the comparison groups shown.

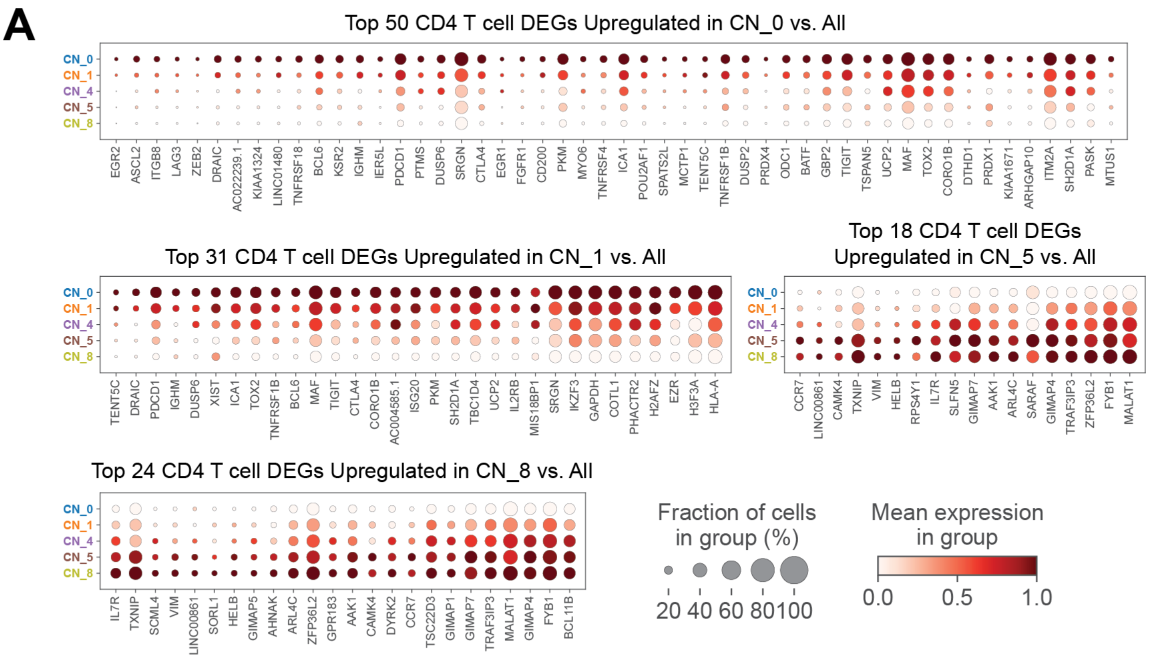


Figure S6: **Additional materials on spatially-resolved differential expression within CD4 T cell populations. (A)** Top upregulated differentially expressed genes (DEGs) and their expression between CD4 T cells from scRNA-seq predicted to originate from a given cell neighborhood compared to all other CD4 T cells. Only cell neighborhoods with ample cell type representation are shown (see cell type composition plot in Figure S3A for more details). Expression for each gene is relative across the comparison groups shown.

#### References


[1] Yuhan Hao, Stephanie Hao, Erica Andersen-Nissen, William M. Mauck, Shiwei Zheng, Andrew Butler, Maddie J. Lee, Aaron J. Wilk, Charlotte Darby, Michael Zager, Paul Hoffman, Marlon Stoeckius, Efthymia Papalexi, Eleni P. Mimitou, Jaison Jain, Avi Srivastava, Tim Stuart, Lamar M. Fleming, Bertrand Yeung, Angela J. Rogers, Juliana M. McElrath, Catherine A. Blish, Raphael Gottardo, Peter Smibert, and Rahul Satija. Integrated analysis of multimodal single-cell data. *Cell*, 2021.

[2] J. Kennedy-Darling, S.S. Bhate, J.W. Hickey, S. Black, G.L. Barlow, G. Vazquez, V.G. Venkataraaman, N. Samusik, Y. Goltsev, C.M. Schürch, and G.P. Nolan. Highly multiplexed tissue imaging using repeated oligonucleotide exchange reaction. *Eur. J. Immunol.*, 2021.

[3] Hamish W. King et al. Single-cell analysis of human b cell maturation predicts how antibody class switching shapes selection dynamics. *Sci Immunol*, 2021.

[4] Hamish W. King et al. Integrated single-cell transcriptomics and epigenomics reveals strong germinal center–associated etiology of autoimmune risk loci. *Sci Immunol*, 2021.

[5] Ville Satopää, Jeannie Albrecht, David Irwin, and Barath Raghavan. Finding a "kneedle" in a haystack: Detecting knee points in system behavior. In *2011 31st International Conference on Distributed Computing Systems Workshops*, pages 166–171. IEEE, 2011.

[6] Adele Cutler and Leo Breiman. Archetypal analysis. *Technometrics*, 36(4):338–347, 1994.

[7] S. He, Y. Jin, A. Nazaret, et al. Starfysh integrates spatial transcriptomic and histologic data to reveal heterogeneous tumor–immune hubs. *Nat Biotechnol*, 2025.

[8] Harold W. Kuhn. The hungarian method for the assignment problem. *Naval Research Logistics Quarterly*, 2(1–2):83–97, 1955.

[9] Andrew Chen, Andy Chow, Aaron Davidson, et al. Developments in mlflow: A system to accelerate the machine learning lifecycle. In *Proceedings of the Fourth International Workshop on Data Management for End-to-End Machine Learning*, pages 1–4, 2020.

[10] S. Chen, B. Zhu, S. Huang, et al. Integration of spatial and single-cell data across modalities with weakly linked features. *Nat Biotechnol*, 2024.

[11] G. Wang, J. Zhao, Y. Lin, et al. scmodal: a general deep learning framework for comprehensive single-cell multi-omics data alignment with feature links. *Nat Commun*, 2025.

[12] Cornelis Joost van Rijsbergen. *Information Retrieval*. Butterworths, London, 2 edition, 1979.

[13] Lawrence Hubert and Phipps Arabie. Comparing partitions. *Journal of Classification*, 2(1):193–218, 1985.

[14] Peter J. Rousseeuw. Silhouettes: A graphical aid to the interpretation and validation of cluster analysis. *Journal of Computational and Applied Mathematics*, 20:53–65, 1987.

[15] Ilya Korsunsky, Nicholas Millard, Jean Fan, Kamil Slowikowski, Fan Zhang, Kevin Wei, Yuriy Baglaenko, Michael Brenner, Po-Ru Loh, and Soumya Raychaudhuri. Fast, sensitive and accurate integration of single-cell data with harmony. *Nature Methods*, 16:1289–1296, 2019. Introduces LISI/iLISI for neighborhood-level modality mixing.

[16] Maren Büttner, Zhichao Miao, F Alexander Wolf, Sarah A Teichmann, and Fabian J Theis. A test metric for assessing single-cell rna-seq batch correction. *Nature Methods*, 16(1):43–49, 2019.

[17] J. Dou, S. Liang, V. Mohanty, et al. Bi-order multimodal integration of single-cell data. *Genome Biol*, 2022.

[18] P. Demetci, R. Santorella, B. Sandstede, et al. Scot: Single-cell multi-omics alignment with optimal transport. *Journal of Computational Biology*, 29(1):3–18, 2022.

[19] K. Cao, X. Bai, Y. Hong, and L. Wan. Unsupervised topological alignment for single-cell multi-omics integration. *Bioinformatics*, 36(Supplement$_1$) $: i48 - -i56, 2020.$

[20] A. Gayoso, Z. Steier, R. Lopez, et al. Joint probabilistic modeling of single-cell multi-omic data with totalvi. *Nat Methods*, 2021.

[21] Domenick E. Kennedy and Marcus R. Clark. Compartments and connections within the germinal center. *Frontiers in Immunology*, 2021.

[22] U. Klein and R. Dalla-Favera. Germinal centres: role in b-cell physiology and malignancy. *Nat Rev Immunol*, 2008.

[23] N. De Silva and U. Klein. Dynamics of b cells in germinal centres. *Nat Rev Immunol*, 2015.

[24] Daisy Y. Ding, Zeyu Tang, Bowen Zhu, et al. Quantitative characterization of tissue states using multiomics and ecological spatial analysis. *Nature Genetics*, 2025.

[25] S. Crotty. Follicular helper cd4 t cells (tfh). *Annu Rev Immunol*, 2011.

[26] CH. Koh, S. Lee, M. Kwak, et al. Cd8 t-cell subsets: heterogeneity, functions, and therapeutic potential. *Exp Mol Med*, 2023.
